# Brain rhythms control microglial response and cytokine expression via NFκB signaling

**DOI:** 10.1101/2023.03.03.530908

**Authors:** Ashley Prichard, Kristie M. Garza, Avni Shridhar, Christopher He, Sara Bitfaran, Yunmiao Wang, Matthew C. Goodson, Dieter Jaeger, Levi B. Wood, Annabelle C. Singer

## Abstract

Microglia, the brain’s primary immune cells, transform in response to changes in sensory or neural activity, like sensory deprivation. However, little is known about how specific frequencies of neural activity, or brain rhythms, impact microglia and cytokine signaling. Using visual noninvasive flickering sensory stimulation (flicker) to induce electrical neural activity at different frequencies, 40Hz, within the gamma band and 20Hz, within the beta band, we discovered these brain rhythms differentially affect microglial morphology and cytokine expression in healthy animals. We found that flicker induced expression of certain cytokines, including IL-10 and M-CSF, that was independent of microglia. Because NFκB is activated by synaptic activity and regulates cytokines, we hypothesized this pathway plays a causal role in frequency-specific cytokine and microglial responses. Indeed, we found that after flicker, phospho-NFκB co-labeled with neurons more than microglia. Furthermore, inhibition of NFκB signaling by a small molecule inhibitor down-regulated flicker-induced cytokine expression and attenuated flicker-induced changes in microglia morphology. These results reveal a new mechanism through which brain rhythms affect brain function by altering microglia morphology and cytokines via NFκB.

**Teaser:** Frequency-specific brain rhythms regulate cytokine expression, microglia morphology, and microglia-independent expression of M-CSF and IL10 via NFκB.

## Introduction

Brain rhythms are found in many brain structures and result from coordinated electrical neural activity at specific frequencies in response to task demands and environmental stimuli (*1–3*). Gamma frequency neural activity (∼30-100Hz) increases during attention and sensory processing, whereas beta frequency (∼12.5-30Hz) is associated with active concentration and suppression of motor behaviors (*2, 4*). Furthermore, neural activity is modulated rhythmically in response to rhythmic sensory stimuli and this entrainment enhances sensory processing, including attention (*5*). While brain rhythms and neural entrainment to sensory stimuli are well-known to reflect and influence neuronal electrical function, little is known about how different frequencies of neural activity impact other brain functions, such as immune signaling (*6*). Our recent work has shown that gamma frequency (40Hz) noninvasive flickering sensory stimulation (flicker) recruits microglia to clear pathogens in mouse models of Alzheimer’s disease, revealing that this specific frequency of neural activity alters microglia, the primary immune cell of the brain in the presence of Alzheimer’s pathology (*7–11*). However, microglia responses vary with pathological context and therefore it is unclear whether these effects of gamma flicker on microglia generalize to healthy brains (*12*). Furthermore, the mechanisms by which gamma frequency activity alters microglia and immune signaling pathways remain unknown.

Brain immune function is typically ascribed to microglia, the primary phagocytes in the brain, because of their roles in engulfing pathogens and secreting both cytokines and related extracellular signals that support neuronal health (*13–15*). Microglia are known to respond to changes in sensory and neural activity, like sensory deprivation or seizures (*16–18*). In the healthy adult brain, microglial processes dynamically shift to survey surrounding synapses, acquiring elongated phenotypes to scan their environment and monitor neuronal activity (e.g., hyper-ramified). Glia rapidly change their phenotype through intermediate stages to shortened processes and enlarged soma when detecting pathogens (e.g., hypo-ramified), and evolving into amoeboid phenotypes that can lack processes altogether (*19*). Nevertheless, little is known about how specific patterns of neural activity, beyond gross increases or decreases, affect microglia and cytokines in healthy individuals. While microglia are a commonly accepted source of cytokines, neurons are known to secrete specific cytokines, like fractalkine (CX3CL1) (*20–23*). Although the mechanisms that govern neuronally-sourced cytokines have received limited attention, cytokine expression is controlled by canonical phospho-protein signaling pathways which are present in both microglia and neurons, including the NFκB and MAPK signaling cascade (*24–27*). Indeed, NFκB is activated in neurons by synaptic activity, and we have shown that 40Hz flicker increases activation of this pathway (*12*). Because of its regulation by synaptic activity and its role in cytokine expression, the NFκB pathway could be a novel mechanism by which brain rhythms regulate microglial responses (*28–31*).

Sensory experiences can induce frequency-specific brain rhythms, like gamma, and neural activity is modulated by rhythmic sensory stimuli. Thus, it is important to understand how specific frequencies of neural activity affect cytokines and microglia. Indeed, cytokine and microglial responses to specific frequencies of neural activity may play a role in how the brain adapts to the environment. Flicker stimulation, which precisely induces specific frequencies of neural activity, allows us to non-invasively test the effects of different frequencies of neural activity without significantly increasing or decreasing mean firing rates, which has more commonly been investigated. We and others have shown that simple flickering lights and sounds, similar to a fast strobe light and beeping, drive specific frequencies of neural activity in primary sensory areas at the flicker frequency of the stimulus (*7–9, 32, 33*) and beyond (*34, 35*). Thus, flicker stimulation non-invasively induces specific frequencies of neural activity to test how this activity affects microglia and cytokines without grossly changing average neural firing rates (*8, 9*). Previously, we have shown that 20Hz and 40Hz visual flicker led to different patterns of cytokine protein expression in healthy mice, but the effects of specific flicker frequencies on microglia in healthy mice and the causal biomolecular mechanisms by which neural activity causes these changes are unknown.

To elucidate how brain rhythms modulate microglia and cytokines, we tested the hypothesis that flicker differentially affects microglia and cytokines via NFκB, which is a canonical phospho-protein signaling pathway. First, we established frequency-specific effects of flicker on microglia and cytokine-related transcription in healthy mice. We used 20Hz and 40Hz flickering stimuli because they drive frequencies of neural activity within common, naturally occurring brain rhythms, and we previously discovered that specific frequencies have distinct effects on cytokine protein levels. We found that 1 hour of 40Hz or 20Hz flicker was sufficient to differentially transform microglia morphology, consistent with different functional roles, and to alter transcription of genes that control cytokine expression (*12*). We next investigated the cellular source of flicker-induced cytokine expression because cytokines can both cause microglial changes or arise from microglia. Importantly, we found that 40Hz flicker in microglial-depleted mice increases tissue levels of the cytokines M-CSF, which recruits microglia, and IL-10, which is anti-inflammatory, indicating that these cytokines are expressed from a different cellular source. Because NFκB and MAPK pathways regulate synapses and immune function, we tested the hypothesis that these pathways are necessary for flicker-induced cytokine expression and microglial changes. We discovered that inhibition of either NFκB or MAPK signaling pathways prior to flicker exposure down-regulated expression of cytokines, while inhibition of NFκB only prevented microglial branching responses to 40Hz flicker. Together, these results show that specific frequencies of flicker elicit a distinct effect on microglia and cytokine transcription. Furthermore, these findings reveal a new NFκB mediated signaling mechanism by which 40Hz flicker activity affects cytokines and microglia. Collectively, our data thus define a previously unknown pathway linking flicker stimulation, brain rhythms, and brain immune signaling.

## Results

### Visual flicker elicits corresponding frequency-specific activity in primary visual cortex

We first established that visual flicker induces neural activity at the specific frequency of flicker across all of visual cortex. While prior work has shown that visual flicker induces neural activity at the specific frequency of flicker in the visual cortex using local field potential recordings (*20, 36*), such recordings do not assess neural activity on a large spatial scale across cortex or even across all of primary visual cortex. Thus, we began the current study by asking if both 20Hz and 40Hz flicker stimuli would drive 20Hz and 40Hz neural activity across visual cortex. To test this, we performed wide-field voltage imaging during 20Hz or 40Hz flicker with a fast voltage indicator, JEDI-1P-Kv, that can detect high frequency neural activity (*36*). We performed widefield fast voltage imaging at 200 fps in awake mice during 20Hz or 40Hz visual flicker to detect fast fluctuations in voltage within neurons. We observed strong neural responses in the frequency of the visual flicker throughout bilateral primary visual cortex at both 20Hz and 40Hz (**Fig. 1B, Fig. S1**). Thus, 20Hz and 40Hz flicker induces concomitant 20Hz and 40Hz neural activity.

**Figure 1.**
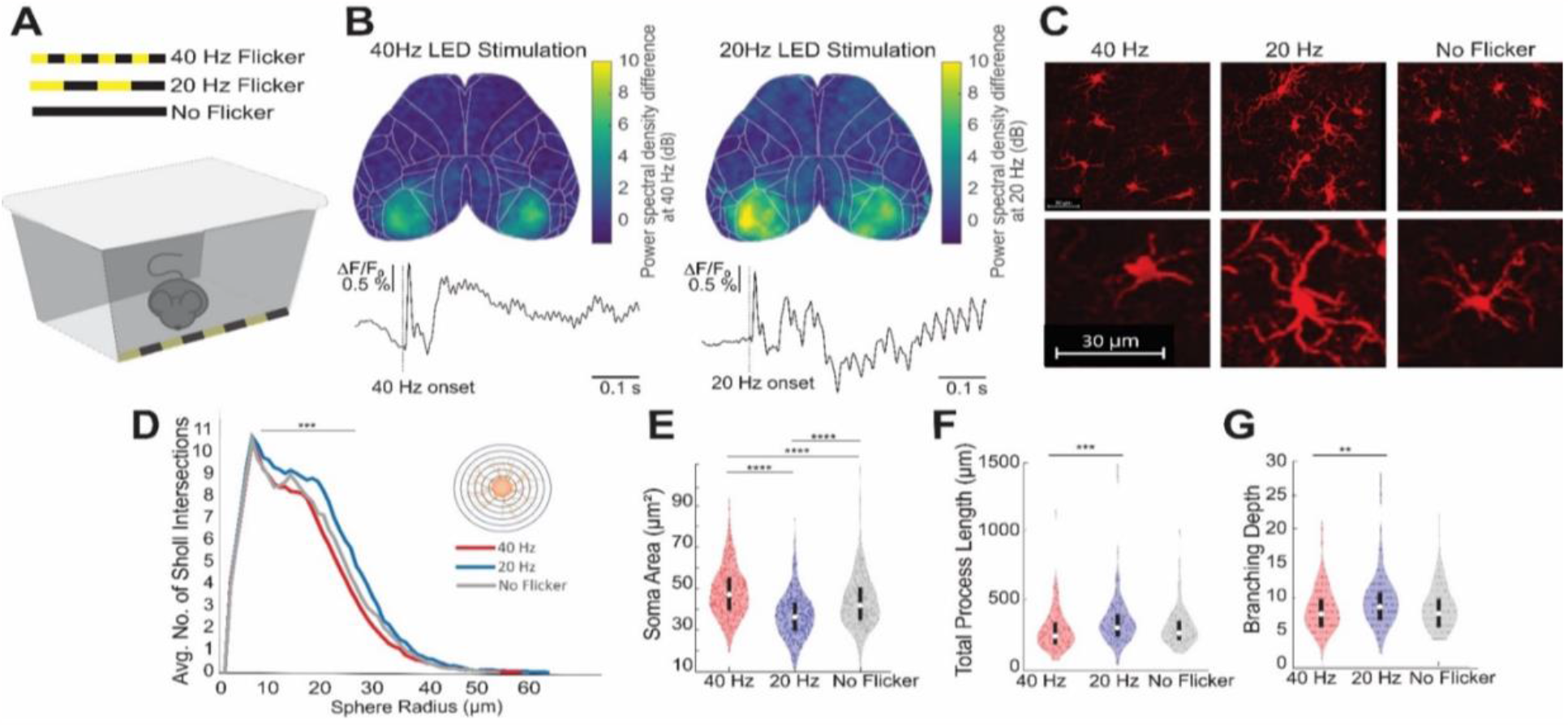
Visual flicker affects microglial morphology in a frequency specific manner in WT mice. **(A)**, Schematic of visual LED-lined stimulation box for mice. **(B), Top:** Spatial heatmap of power differences at 40Hz (left) or 20Hz (right) during visual flicker relative to no flicker from widefield voltage imaging of JEDI-1P-Kv from 1 representative mouse. Both spatial maps share the same scale bar determined by the maximum and minimum values across the two sets of data for visualization. **Bottom:** Mean ΔF/F_0_ trace from 2×2-pixel (200 μm × 200 μm) at primary visual cortex indicating the oscillatory response during 40Hz (left) or 20Hz (right) visual flicker. Vertical line indicates flicker onset. n = 6 mice and 60 trials, 10 trials/mouse. **(C)**, Example images of IBA-1+ microglia (red) from each experimental condition show clusters of representative microglia (top row, taken at 20x) and a single representative microglia (bottom row, taken at 40x). Scale bar indicates 30 μm. (**D)**, Sholl analysis of IBA-1+ microglia after animals were exposed to 1hour of 40 Hz Flicker (red, n=6 mice, 174 microglia), 20 Hz Flicker (blue, n=5 mice, 140 microglia), or No Flicker (grey, n=5 mice, 132 microglia) shows significant differences in the number of branches between flicker groups (linear mixed model with repeated measures; F (2,15271) = 17.65, p < 0.001), and at different radii extending from the soma (F(1,15271) = 3034.051, p < 0.001). These differences between groups occurred at specific radii: significant differences between 40Hz, 20Hz, and No Flicker on the number of branches occurred within 30 μm of the cell body (main effects for intercept and frequency at bins 0-10 (F(1,4897)=12325.18, p <0.001), (F(2,4897) = 4.84, p =0.008); bins 11-20 (F(1,4423)=13025.2, p <0.001), (F(2,4423) = 38.17, p <0.001); and bins 21-30 (F(1,3809)=5328.96, p <0.001), (F(2,3809) = 39.638, p <0.001), but were not significant at higher radii. (**E)**, Soma area of IBA-1+ microglia differed after 40Hz (n=6 mice, 459 microglia), 20Hz (n=5 mice, 398 microglia), or No Flicker (n=5 mice, 388 microglia) (F(12,1242)=98.634467 p=1.75E-40; post-hoc t-tests: 40Hz vs. 20Hz: p = 9.56E-10, 40Hz vs No Flicker: p = 1.04E-8, 20Hz vs No Flicker: p = 9.56E-10; Tukey’s HSD corrected for multiple comparisons). **(F)**, Total process length of IBA-1+ microglia differed after 40Hz or 20Hz flicker (F(2,443)=6.8925, p=0.0011; post-hoc t-tests: 40Hz vs. 20Hz: p = 0.0007, 40Hz vs No Flicker: p = 0.432, 20Hz vs No Flicker: p = 0.0622; Tukey’s HSD corrected for multiple comparisons). **(G)**, Full branching depth of microglia differed after 40Hz or 20Hz flicker (F(2,443)=4.789, p=0.0088; post-hoc t-tests: 40Hz vs. 20Hz: p = 0.0069, 40Hz vs No Flicker: p = 0.7235, 20Hz vs No Flicker: p = 0.0915; Tukey’s HSD corrected for multiple comparisons). For **(E-G)**, F- and p-values were generated from one-way, two-tailed, unpaired ANOVA tests. Differences between groups were found from one-way, two-tailed, t-tests with Tukey’s HSD correction. Box plots inside violin plots indicate median and quartiles, dots indicate individual microglia. **p<0.01,***p<0.001,****p<0.0001.

### Visual 20Hz and 40Hz flicker stimuli elicit distinct microglial morphological responses in healthy mice

Having established that 20Hz and 40Hz flicker induce different brain rhythms throughout primary visual cortex, we next asked if 20Hz and 40Hz stimulation led to different microglial morphologies that correspond to different microglial functions (*19, 37*), *12, 38*). We hypothesized that 40Hz flicker would induce a transition to hypo-ramified microglial morphology, as observed during synaptic engulfment and other microglia activity, while 20Hz flicker would induce microglia with hyper-ramified morphology that occurs during surveillance or homeostatic processes (*14, 38*). We expected that while 20Hz and 40Hz would elicit different microglia responses, these effects should be more subtle than those due to injury or disease, instead reflecting a range of microglia morphology present in the healthy brain (*37, 39–42*). To test this hypothesis, we analyzed microglial morphology after the mice were exposed to 1hour of flicker at either 20Hz flicker, 40Hz flicker, or no flicker (constant light) (**Fig. 1A,C**). We found that the 40Hz flicker elicited a significant increase in soma area compared to 20 Hz whereas 20Hz elicited significantly increased process length and branching compared to 40Hz (**Fig. 1E-G, Fig. S2**). However, these changes were not associated with significant differences in Iba1 (encoded by *Aif1*) transcription (p=0.6 unadjusted, DESeq2). Thus, 40Hz stimulation leads to more amoeboidal morphology with less ramification consistent with engulfing microglial phenotypes, consistent with ours and other prior studies (*19, 37, 43*) and 20Hz stimulation results in a more ramified morphology with longer processes consistent with surveillance phenotypes.

### RNAseq analysis reveals differences in cytokine-related transcription factors between 20Hz and 40Hz flicker

To identify potential molecular mechanisms of these different microglia responses to brain rhythms, we asked which immune-related gene pathways also differentially respond to 20Hz and 40Hz flicker. To broadly examine changes in expression of several potentially microglia-related pathways with relatively high sequencing depth, we used bulk RNAseq to quantify gene expression in the visual cortices of mice exposed to 20Hz or 40Hz combined audio/visual flicker, which is expected to have similar effects in the visual cortex as visual-only flicker (*12*). We then used a gene set variation analysis (GSVA) with the C2 canonical signaling pathways gene sets database (MSigDB, Broad) (*44*). We found different frequencies of flicker differentially affect transcription factors that control cytokine expression with a more nuanced effect than injury or disease. Interestingly, we did not identify clear changes in signaling pathways that have previously been shown to affect microglial response to neural activity, including norepinephrine and chemoreceptor P2Y12 (**Fig. S3**). Importantly, we also did not identify any significantly enriched gene sets between 20Hz and 40Hz flicker related to immunity, including interleukins pathway, MAPK pathway, cytokine signaling, stress response, or complement cascade (**Fig. 2A, Table 1**). However, these cytokine-related immune pathways are made up of broad lists of more than 50 genes, and a more detailed examination revealed significant differences in expression of transcription factors that control cytokine expression, such as Ifit1 and Irak4 (*45–47*) (**Fig. 2B-C**). While the 20Hz and 40Hz flickers induced significant differences in cytokine expression, neither produced coordinated expression patterns indicative of broad inflammation that is often found in LPS models, for example (*48*). Given that these changes in cytokine expression are associated with changes in microglial morphology and occur in healthy brains, they suggest an overall frequency-specific modulation of cytokines that is distinct from that due to injury or disease.

**Figure 2.**
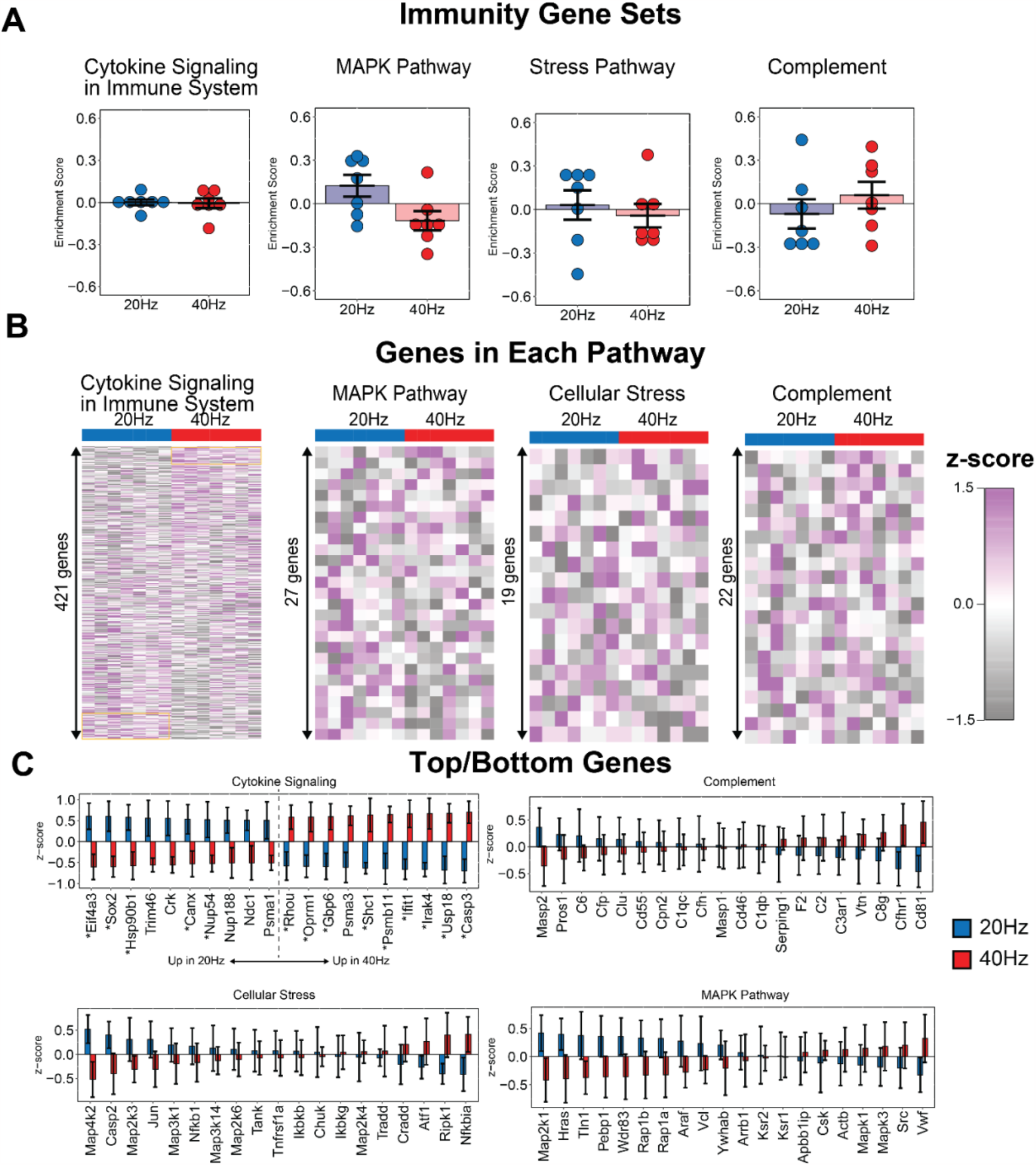
Audio/visual flicker for 1 hour at 40 Hz significantly down-regulates divergent expression of cytokine pathways. **(A)**, Gene set variation analysis did not identify significant enrichment immune-related gene sets after 20Hz (blue) or 40Hz (red) flicker (mean±SEM, statistical testing via gene set permutation analysis, dots indicate individual animals). **(B)**, Heatmap representation of all genes within each pathway reveals that 20 vs 40Hz flicker drive distinct patterns of gene expression within the ‘Cytokine Signaling’ pathways but no clear patterns in other pathways (rows are z-scored, each row is a gene, each column is an animal). (**C**), All of the top and bottom 10 genes within the ‘Cytokine Signaling’ pathway are each significantly differentially regulated (top left, DESeq2, p<0.05 un-adjusted). Top/bottom 10 genes for ‘Complement’ (top right), ‘Cellular Stress’ (bottom left), and ‘MAPK Pathway’ (bottom right) signaling show no significant differences.

### M-CSF and IL-10 protein expression is independent of microglia

Because cytokines both signal to and are secreted by microglia, we determined the necessity of microglia for flicker-induced cytokine expression by examining protein expression changes after depleting microglia. We compared 40Hz effects to those of 20Hz because these stimuli differentially affected cytokines and microglia and because 20Hz flicker controls for key aspects of the flickering stimulus, including light exposure, stimuli turning on and off repeatedly, and any salience associated with flickering stimuli. We depleted microglia using PLX 3397 (290ppm in the chow, **Fig. S4** see **Materials and Methods**), exposed mice to visual flicker, and then used a Luminex multiplexed immunoassay to quantify protein expression of 32 cytokines in the visual cortex, including many that previously showed differential expression in response to 40Hz and 20Hz flicker in our prior work (**Fig. 3**)(*12*). We initially compared protein expression in control animals fed a regular diet versus PLX diet, showing that expression of some cytokines persisted in microglia-depleted mice (**Fig. S5**). We accounted for the multi-dimensional nature of the data using a discriminant partial least squares regression (PLSDA), as we have done previously (*12*), We identified two latent variables to separate flicker frequency condition (latent variable 1, LV1) and intact versus depletion of microglia (latent variable 2, LV2). LV1 separated 40Hz flickered animals to the right and 20Hz flickered animals to the left along latent variable 1 (LV1, **Fig. 3B**). Additionally, microglia depleted animals (40Hz PLX) separated toward the top from microglial-intact animals toward the bottom, along LV2. We evaluated potential overfitting of the model using a permutation analysis wherein sample labels were permuted 1000 times, and distances between group centroids were compared against true group labels. This analysis revealed that true group assignments were better than random (40Hz vs 20Hz: p=0.316, 40Hz vs 40Hz PLX: p=0.20, 40Hz PLX vs 20Hz: p=0.107).

**Figure 3.**
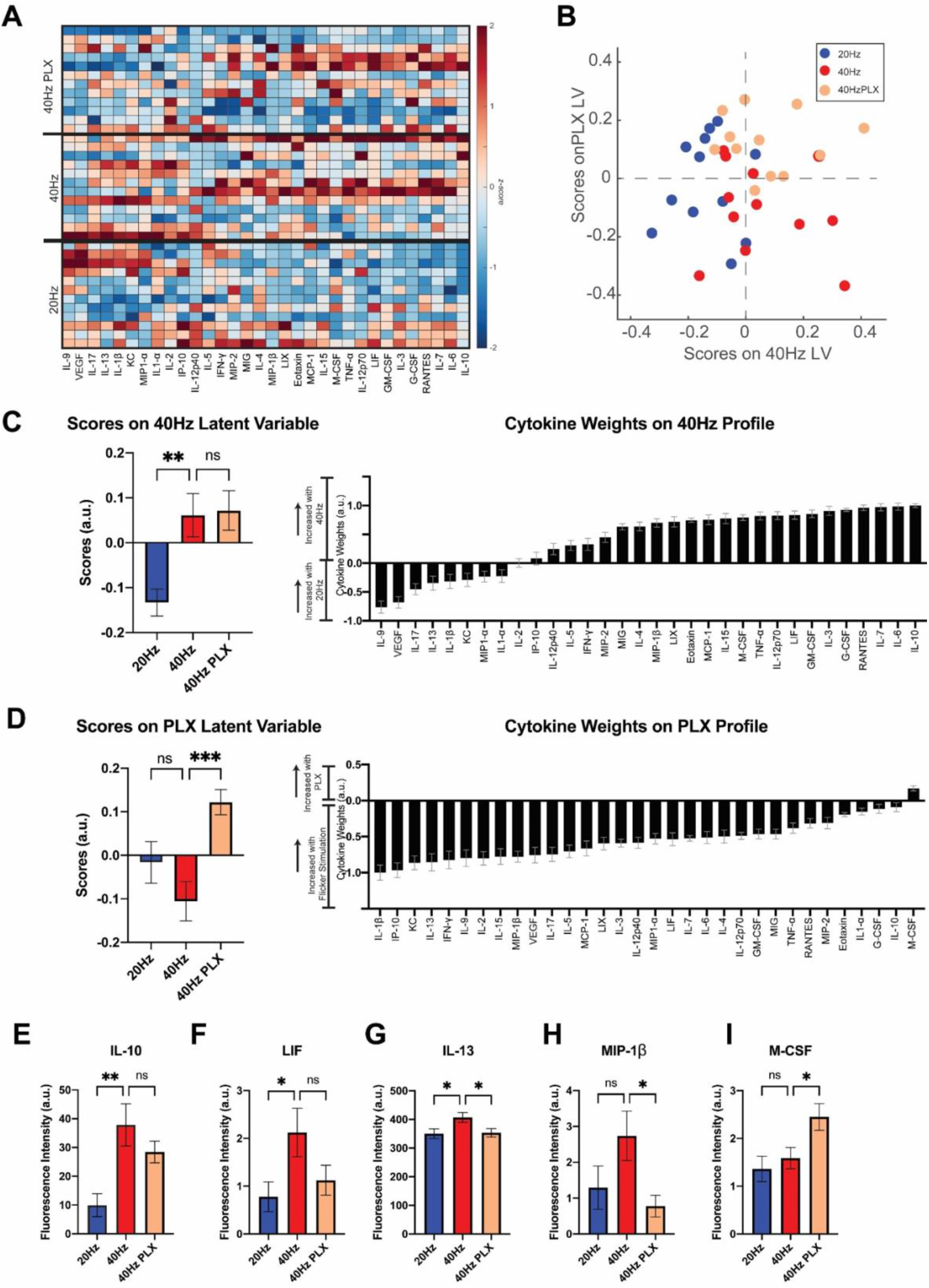
40Hz flicker upregulates expression of cytokines MCSF and IL10 in microglial-depleted mice. **(A)**, Cytokine expression in visual cortices of mice fed PLX and exposed to 1hour of 40Hz flicker (top section “40Hz PLX”) or control diet then exposed to 40Hz (center “40Hz”) or 20Hz (bottom “20Hz”) flicker. Cytokines (columns) are arranged in the order of their weights on the LV1. Color indicates z-scored expression levels for each cytokine. Each row is an animal. **(B)**, PLSDA identified LV1, the axis that separated 40Hz flicker-exposed animals (red and beige) to the right and 20Hz flicker (blue)-exposed animals to the left.LV2 separated animals treated with PLX and 40Hz flicker (beige) on top versus control and 40Hz flicker exposure (red) on the bottom. **(C)**, left, LV1 (termed 40Hz Latent Variable) scores were significantly different between groups (mean ± SEM; F(2,33)=7.747, p=0.0017, one-way ANOVA) and post-hoc analysis revealed significant differences between animals treated with control diet and 40Hz flicker versus 20Hz flicker (p=0.0043, Dunnett’s multiple comparisons). Right, the weighted profile of cytokines that make up LV1 or 40Hz Profile based on which cytokines best correlated with separation of 40Hz (positive) versus 20Hz flicker control (negative) (mean ± SD from a leave-one-out cross-validation). **(D), Left**, LV2 (termed PLX Latent Variable) scores were significantly different between groups (mean ± SEM; F(2,33)=7.630, p=0.0019, one-way ANOVA) and post-hoc analysis revealed significant differences between animals exposed to 40Hz flicker and pretreated with PLX diet or control diet (p=0.0010, Dunnett’s multiple comparisons). **Right**, the weighted profile of cytokines that make up LV2, or PLX Profile, based on which cytokines best correlated with separation of animals treated with PLX (positive) versus flicker (negative) (mean ± SD from a leave-one-out cross-validation). **(E)**, Expression of IL-10 differed across the three groups (mean ± SEM; F(2,33)=7.195, p=0.0026, one way ANOVA), with a significant increase in expression in animals exposed to 40Hz control versus 20Hz control (mean ± SEM; IL-10: p=0.0014). (**F)**, Expression of LIF differed between the 40Hz and 20Hz control groups (p=0.0362). **(G)**, Expression of IL-13 differed across groups (mean ± SEM; F(2,33)=3.840, p=0.0317, one-way ANOVA). A post-hoc comparison test showed significant differences between 20Hz and 40Hz flicker exposed animals (p=0.0345) and between 40Hz flicker control and PLX pretreated animals (p=0.0479). **(H)**, Expression of MIP-1β differed across groups (mean ± SEM; F(2,33)=3.315, p=0.0488, one-way ANOVA) and post-hoc tests show significant differences between the 40Hz control group and 40Hz plus PLX diet (p=0.0339). **(I)**, Expression of M-CSF differed across groups (mean ± SEM; F(2,33)=4.959, p=0.0131, one-way ANOVA). Post-hoc multiple comparisons analysis shows significant differences between animals exposed to 40Hz and 40Hz plus PLX diet (p=0.0445).

We identified which cytokines had decreased or intact expression levels when microglia were depleted. First, we determined the profile of cytokines in LV1 (**Fig. 3C**) that were associated with 40Hz (positive) or 20Hz (negative) and the profile of cytokines in LV2 (**Fig. 3D)** that were associated with microglial depletion (PLX, positive) vs. microglial intact mice (negative). Interestingly, top correlates from LV1, including IL-10 which has anti-inflammatory functions, were not significantly changed by microglial depletion, showing it is microglia-independent (**Fig. 3E**). In contrast, the top correlate from LV2, was M-CSF (CSF1, the agonist for CSF1R, which is antagonized by PLX3397), which was increased by microglial depletion and is known to recruit microglia, showing that its expression levels were microglia-dependent but lead to increased rather than decreased expression (**Fig. 3I**). Importantly, some cytokines including IL-13, an anti-inflammatory cytokine, and MIP-1β, a chemokine, were microglial-dependent, consistent with the known role of microglia in cytokine expression (*49*). Surprisingly, flicker-induced expression of IL-10 and M-CSF is independent of microglia or increased in the absence of microglia. Thus, flicker stimulation increased expression of both microglial-dependent and -independent cytokines.

### Phosphorylated NFκB colocalizes with neurons following flicker

We hypothesized that the increased expression of flicker-induced cytokines may arise from NFκB signaling because this pathway is activated in neurons by synaptic activity and controls cytokine expression (*28–31*). Indeed, we previously found increased phosphorylation of the NFκB pathway in visual cortex after 15 min of 40Hz flicker (*12*). To test our hypothesis, we first asked if NFκB is activated in neurons or microglia by 40Hz flicker using immunohistochemistry to co-label phosphorylated NFκB (pNFκB) with cell markers for neurons (NeuN) and microglia (Iba1). We examined co-labeling after 15 min of flicker because phosphorylation of this pathway is transient and was significantly activated after 15 min of flicker in our prior work (*12*). After 40Hz flicker stimulation, pNFκB colocalized with the majority of NeuN+ labeling and little Iba1 labeling. Specifically, 43% (SEM=0.064) of pNFκB signal colocalized with neurons compared to 22% (SEM=0.04) of microglia (F(1,26)=8.95, p = 0.006) (**Fig. 4**). These findings indicate that NFκB signaling occurs within neurons after flicker. There was a significant effect of flicker condition (F(2,26) = 5.56, p=0.01), however Tukey’s HSD post-hoc tests revealed that the effect of flicker condition was due to significant differences between the flicker conditions 40Hz vs Light for NeuN (p=0.008).

**Figure 4.**
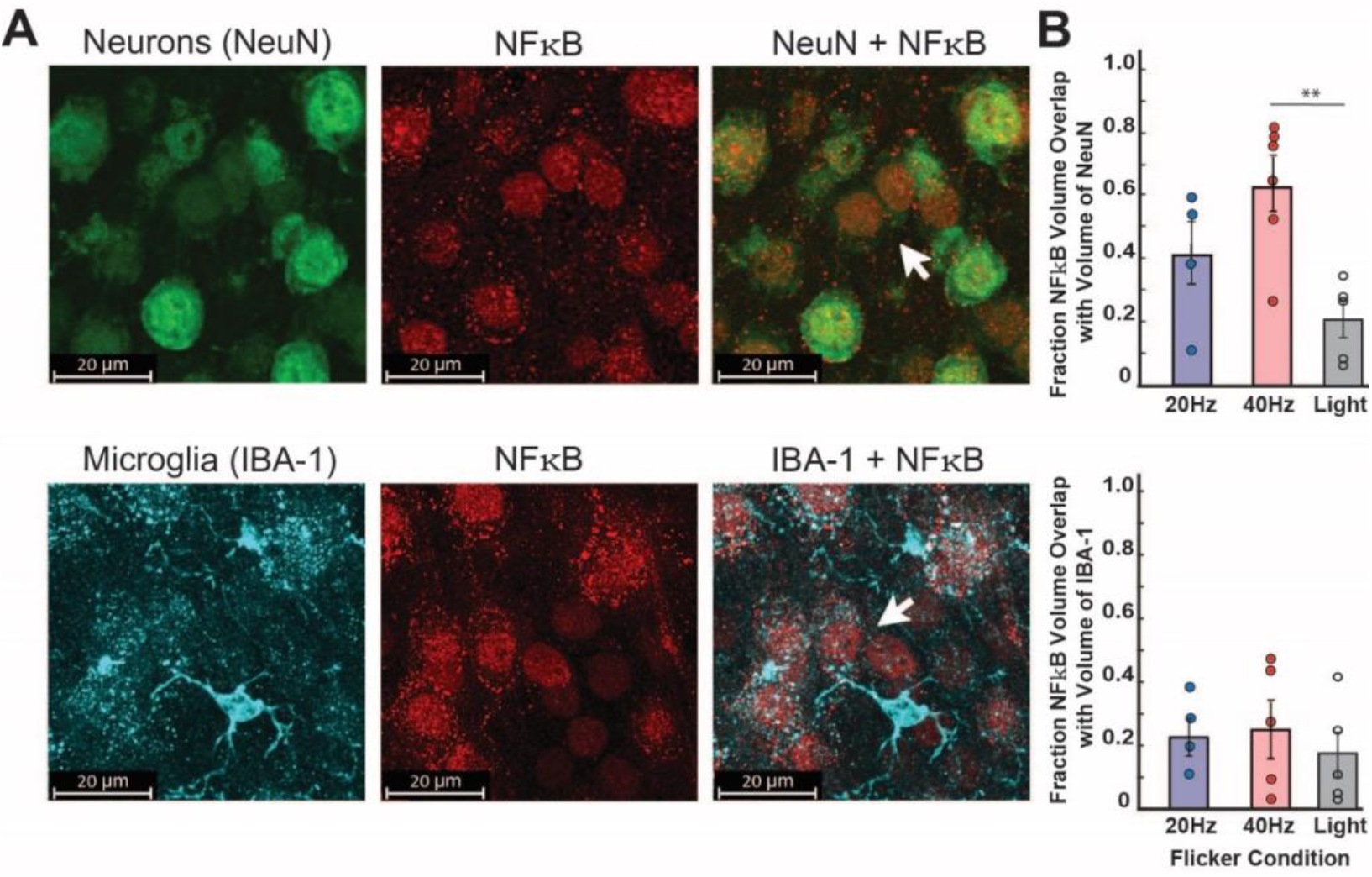
pNFκB colocalizes with neurons after exposure to 40Hz flicker. **(A) Top**, Representative images show co-localization of pNFκB volume (red) with neuronal marker, NeuN, (green) volume after 15 minutes of 40Hz flicker exposure (scale bar= 20 μm); **Bottom**, Representative images labeling microglia marker IBA-1, (cyan), shows low colocalization of pNFκB (red) volume within microglia volume after 15 minutes of 40Hz flicker exposure (scale bar= 20 μm). **(B)**, Percent of total pNFκB volume colocalized with cells (**Top**, neurons; **Bottom**, microglia) across flicker frequencies. ANOVA results show pNFκB was highly colocalized with neurons (mean=43%, SEM=0.064) compared to microglia (mean=22%, SEM=0.04) (F(1,26) = 8.95, p = 0.006), with a significant effect of flicker condition (F(2,26) = 5.56, p=0.01). Tukey’s HSD post-hoc tests revealed that the effect of flicker condition was due to significant differences between flicker conditions 40Hz vs Light for NeuN (p=0.008) compared to 40Hz vs 20Hz (p=0.17) or 20Hz vs Light (p=0.28).

### Gamma flicker-induced microglial changes are mediated by NFκB

Having found that pNFκB co-labeled with neurons and that microglia-modulating cytokine expression was stimulated by 40Hz flicker in microglial-depleted mice, we next reasoned that that inhibition of predominantly neuronal pNFκB would inhibit changes in microglial morphology. To test this, we examined inhibition of two pathways previously shown to be altered by flicker (*12*). We expanded our approach to examine both NFκB, which localized to neurons, and the MAPK pathway because we previously found that phosphorylation of both pathways are altered by 40Hz flicker and both are known to respond to synaptic activity (*12, 50, 51*). As above, we compared microglial morphology in mice exposed to 40Hz and vehicle, 40Hz and MAPK inhibitors, or 40Hz and NFκB inhibitors. Mice were injected intraperitonealyl with small molecule inhibitors of NFκB (10mg/kg IMD0354 plus 100mg/kg Curcumin), which inhibits phosphorylation of NFκB and its translocation into the nucleus or MAPK (100mg/kg of the MAPK/MEK inhibitor SL327 plus 15mg/kg of the MAPK/JNK inhibitor SP600125) pathways 1 hour prior to exposing animals to flicker, using multiple inhibitors for each pathway because of the known compensatory effects within each pathway. Following exposure to 40Hz flicker, we found that microglia soma area was significantly smaller after NFκB inhibition than MAPK inhibition (pJNK+pERK inhibitor), but were not different from the 40Hz+vehicle control, suggesting these inhibitors have distinct, but mild effects on soma size (F(2,370)=7.5909, p=0.0006; post-hoc Tukey’s HSD: VEH+40Hz vs. MAPK Inhibitor+40 Hz: p = 0.1757, MAPK Inhibitor+40 Hz vs NFkB Inhibitor+40 Hz: p = 0.0795, MAPK Inhibitor+40 Hz vs NFkB Inhibitor+40 Hz: p = 0.0003; Tukey’s HSD corrected for multiple comparisons) (**Fig. 5C**). Assessing branching depth, NFκB inhibition increased process branching when compared to both the 40Hz vehicle group and MAPK inhibitor (F(2,229)=7.3418, p=0.0008; post-hoc t-tests: VEH+40Hz vs. MAPK Inhibitor+40 Hz: p = 0.7765, MAPK Inhibitor+40 Hz vs NFkB Inhibitor+40 Hz: p = 0.0006, MAPK Inhibitor+40 Hz vs NFkB Inhibitor+40 Hz: p = 0.0051; Tukey’s HSD corrected for multiple comparisons) (**Fig. 5D, Fig. S6**). Together, these results demonstrate that microglial process complexity following 40Hz flicker is increased when NFκB is inhibited, showing that NFκB is a required for 40Hz flicker induced decreases in microglia branching, but not increases in cell body diameter. Indeed, NFκB inhibition with 40Hz flicker appears to shift microglia into a more ramified state similar to that elicited by 20Hz flicker (i.e. longer branching depth and process length) rather than the hypo-ramified pattern we observed following exposure to 40Hz flicker, indicating NFκB’s role as a mediator in frequency-specific effects on microglia morphology.

**Figure 5.**
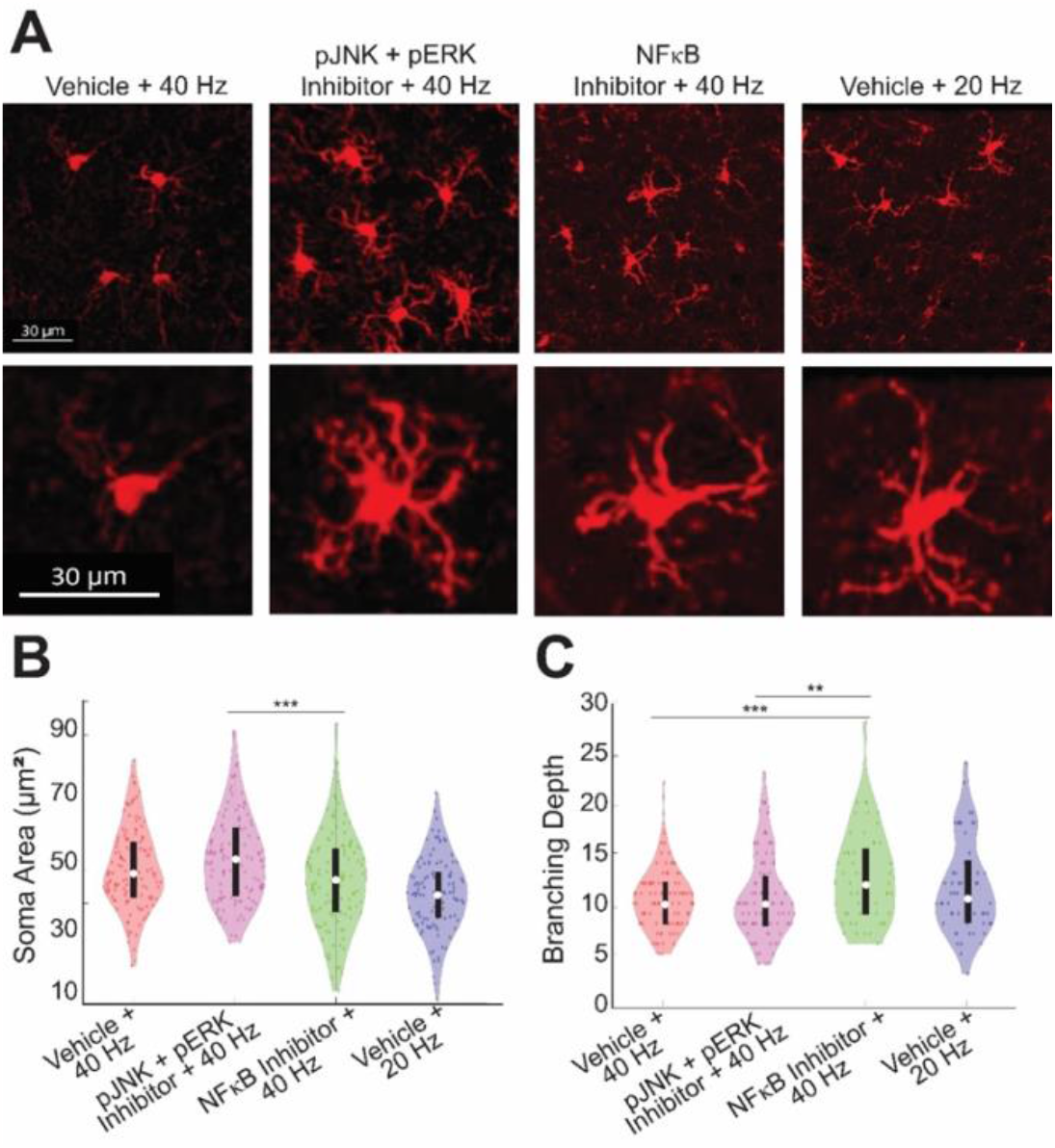
40Hz flicker-induced microglia changes are mediated by phospho-signaling pathways. **(A)**, Example images show clusters of representative IBA-1+ microglia and single representative microglia (bottom) from each experimental condition (Vehicle+40Hz, MAPK Inhibitor (pJNK+pERK inhibitor)+40Hz, NFκB Inhibitor+40 Hz, and Vehicle+20Hz) taken at 40x magnification. Scale bar indicates 30 μm. **(B)**, Soma area of IBA-1+ microglia differed between groups (F(3,475)=14.8797, p=2.84E-9), post-hoc t-tests: MAPK Inhibitor+40 Hz vs NFkB Inhibitor+40 Hz: p = 0.0003, Tukey’s HSD corrected for multiple comparisons, and **(C)**, Full branching depth of IBA-1+ microglia differed between groups (F(2,229)=7.3419, p=0.0008; post-hoc t-tests: VEH+40Hz vs. MAPK Inhibitor+40 Hz: p = 0.7765, MAPK Inhibitor±-40 Hz vs NFkB Inhibitor+40 Hz: p = 0.0007, MAPK Inhibitor+40 Hz vs NFkB Inhibitor+40 Hz: p = 0.0051; Tukey’s HSD corrected for multiple comparisons). For (**B**,**C**) violin plots, F- and p-values were generated from one-way, two-tailed, unpaired ANOVA test. Differences between groups were found from post-hoc Tukey’s HSD multiple-comparisons test. Box plots inside violin plots indicate median and quartiles, dots indicate individual microglia. **p<0.01,***p<0.001,****p<0.0001.

### Cytokine expression is down-regulated by MAPK and NFκB inhibitors

Having found that the NFκB and MAPK signaling pathways mediate flicker-induced changes in microglial morphology, we concluded by asking if inhibition of these pathways affects cytokine expression after 40Hz flicker. To test this, we quantified 32 cytokines using Luminex analysis in mice exposed to 40Hz and MAPK inhibitors, 40Hz and NFκB inhibitors, 40Hz and vehicle, or 20Hz and vehicle. PLSDA revealed that 40Hz samples separated to the right, while 20Hz or 40Hz with either MAPK or NFκB separated to the left along LV1 (**Fig. 6B**). LV1 (**Fig. 6C-D)** consisted of a profile of cytokines associated with 40Hz stimulation, including TNF-α, IL-1β, MIG, IL-15, KC, all of which have pro-inflammatory properties (*51–53*), indicating that inhibition of each pathway suppresses the pro-inflammatory effects of 40Hz compared to 20Hz. The PLSDA also revealed that MAPK inhibition separates toward the bottom and NFκB separates toward the top, along LV2, indicating that each inhibitor has distinct effects on cytokine expression (**Fig. 6E**). As above, the model was evaluated for potential overfitting using a permutation analysis with sample labels permuted 1000 times, and distances between group centroids were compared against true group labels. This analysis revealed that true group assignments were better than random (p_perm_= 1.0 for 40Hz vs 20Hz, 0.128 for MAPK inhibitor vs 40Hz, 0.417 for NFκB inhibitor vs 40Hz, 0.227 for MAPK inhibitor vs NFκB inhibitor, 0.67 for NFκB inhibitor vs 20Hz). Importantly, several of the cytokines we found that were expressed after microglia depletion were decreased following inhibition of NFκB and MAPK including IL-10, MCSF, and LIF (**Fig. S7**), indicating that expression of these cytokines is both independent of microglia and dependent on these phosphor-signaling pathways.

**Figure 6.**
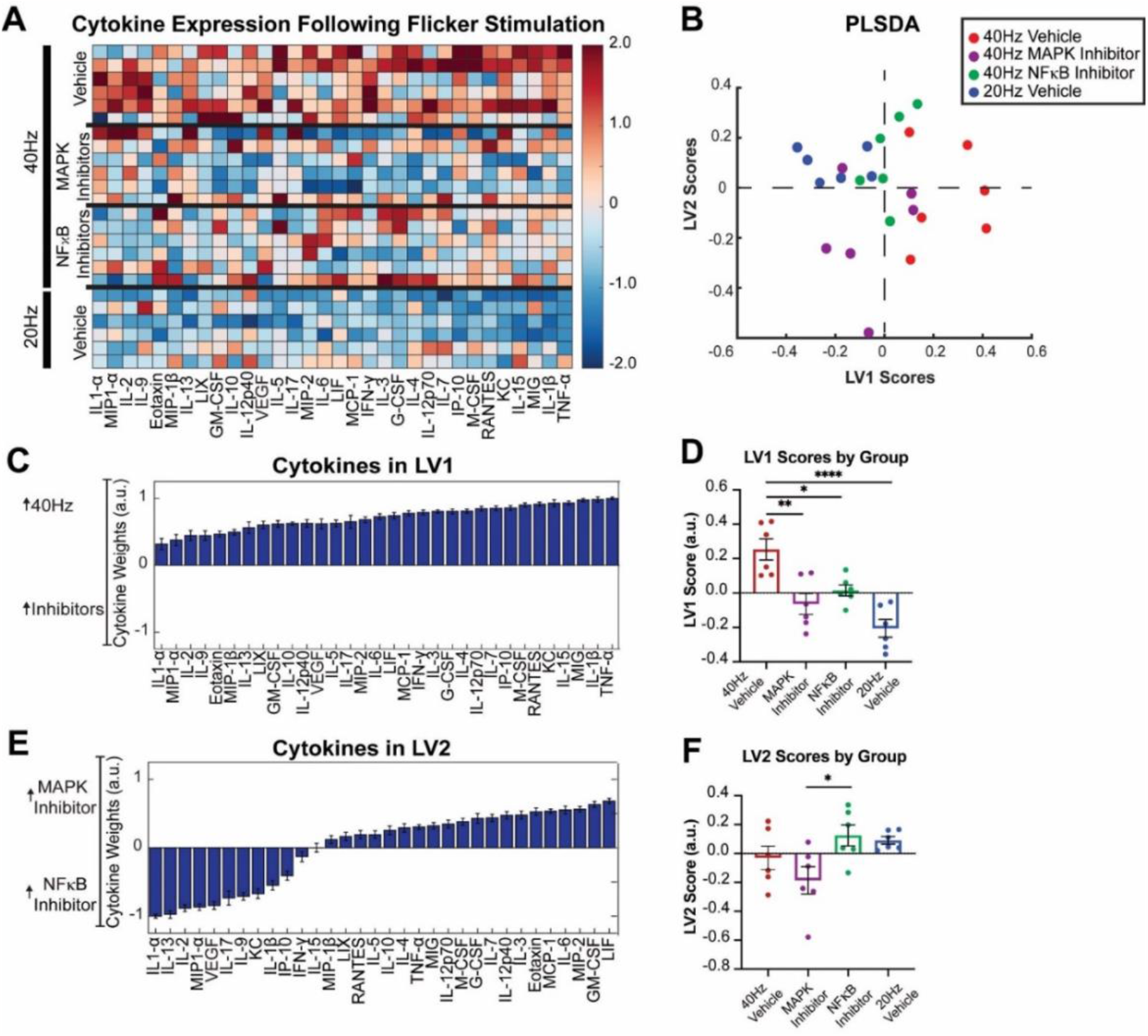
40Hz flicker-induced cytokine protein expression is mediated by MAPK and NFκB pathways. **(A)**, Cytokine expression in visual cortices of mice administered either vehicle (top and bottom), MAPK inhibitors (top center) or NFκB inhibitors (bottom center) phosphosignaling inhibitors and then exposed to 1 hour of 40Hz (top) or 20Hz (bottom) visual flicker. Each row represents one animal (n = 6). Cytokines (columns) are arranged in the order of their weights on the LV1 in **(D)**. Color indicates z-scored expression levels for each cytokine. **(B)**, PLSDA identified LV1, the axis that separated vehicle + 40Hz flicker-exposed animals (red) to the right and all other groups to the left (vehicle + 20Hz flicker, blue; MAPK inhibited + 40Hz flicker, purple; and NFκB inhibited + 40Hz flicker, green). Dots indicate individual animals for all graphs in this figure. LV2 separated the two inhibitor types (NFκB inhibited, green; MAPK inhibited, purple). **(C)**, The weighted profile of cytokines that make up LV1 based on which cytokines best correlated with separation of 40Hz (positive) versus other groups (mean ± SD from a leave-one-out cross-validation). **(D)**, LV1 scores were significantly different for the 40Hz group compared with the comparison groups (mean ± SEM; F(3,20) = 13.28, p < 0.0001, one-way ANOVA). **(E)**, The weighted profile of cytokines that make up LV2 based on which cytokines best correlated with separation of the two inhibited groups MAPK inhibitor (positive) and NFκB inhibitor (negative) (mean ± SD from a leave-one-out cross-validation). **(F)**, LV2 scores were significantly different between the four groups (mean ± SEM; F(3,20) = 3.689, p = 0.0291, one-way ANOVA), with post-hoc analysis revealing a significant difference between animals injected with the MAPK versus NFκB inhibitors (p=0.0415).

In sum, our data show that different frequencies of flicker elicit different brain rhythms with divergent microglial and cytokine responses. Furthermore, our results show that much of the 40Hz flicker induced expression of cytokines arise from microglia, though not all, and that changes in microglia and cytokine expression depend on pNFκB. Our results are thus consistent with a model in which 40Hz flicker induces phosphorylation of NFκB that in turns leads to expression of cytokines that affect microglial phenotypes. The initial effect on microglia may results from microglia-independent cytokines, such as M-CSF and IL-10, that arise from neuronal pNFκB (**Fig. 7**). This in turn could initiate further release of cytokines from microglia. Our results point to novel mechanisms of brain neuron-microglia communication in response to flicker and flicker-induced brain rhythms.

**Figure 7.**
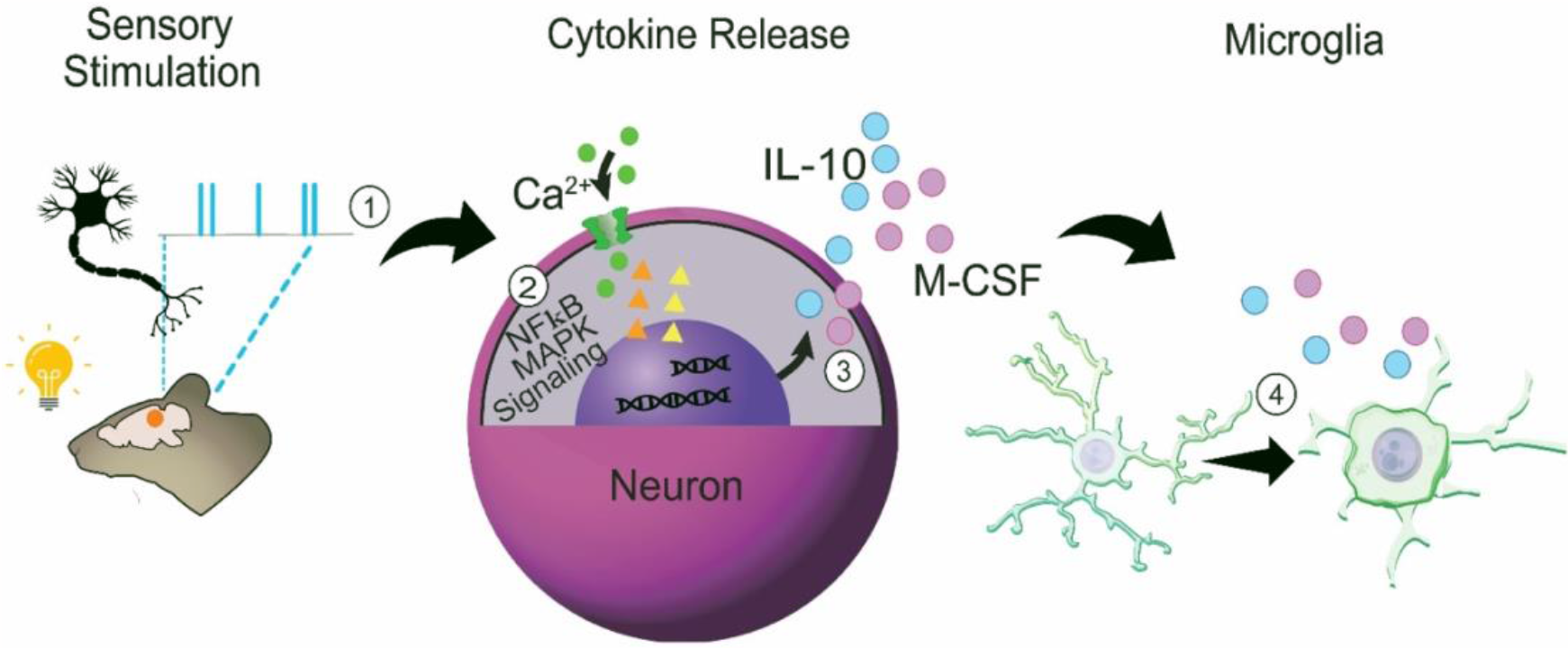
Proposed mechanisms by which 40Hz flicker changes microglia and cytokine signaling. **(1)** Visual flicker entrains neurons in mouse visual cortex, **(2)** activating MAPK and NFκB pathway signaling in the neuron to regulate distinct **(3)** cytokines and subsequently, **(4)** microglial morphology. **(2)** pNFκB labeling was highly colocalized with neurons, more so than microglia, and was present even in the absence of microglia, suggesting that neurons play a key role in the modulation of the neuroimmune environment via NFκB signaling. **(3)** 40Hz flicker increased the expression of several key cytokines including M-CSF, known to recruit microglia, and IL-10, a powerful anti-inflammatory, even in the absence of microglia. **(4)** Frequency-specific changes were evident in microglia morphology accompanying the altered expression of anti-inflammatory cytokines: 20Hz flicker elicited a surveillance morphology phenotype whereas 40Hz flicker elicited an engulfment/ameboid morphology phenotype, suggesting that specific frequencies play an important role in regulating brain immunity. When the NFκB pathway was inhibited 40Hz flicker did not fully induce this ameboid-like state. Furthermore, cytokine expression following flicker was mediated by NFκB phospho-signaling, suggesting that NFκB signaling mediates flicker’s effects on microglial cell morphology and cytokine signaling.

## Discussion

In the current study, we discovered that flicker-induced brain rhythms have frequency-specific effects on microglia and cytokines via activation of NFκB. We found that one hour of visual stimulation at specific frequencies is sufficient to alter microglia morphology into different phenotypes consistent with performing disparate functions (*37*). 40Hz flicker induced 40Hz neural activity, microglia morphology associated with engulfment, and an altered profile of cytokines expression. In contrast, 20Hz flicker induced 20Hz neural activity, microglia surveillance or homeostatic phenotypes, and a different pattern cytokine expression compared to 40Hz. Through microglia depletion, we identified 40Hz flicker-induced cytokines that were microglia-independent, including M-CSF, a microglial chemokine and colony stimulating factor, and IL-10, an anti-inflammatory cytokine, which may in turn induce microglial responses. We then investigated the role of NFκB and MAPK signaling in mediating flicker-induced microglia and cytokine changes because these pathways are both regulated by cytokine expression and responds to synaptic activity (*28–31*). Importantly, pNFκB was highly localized within neurons compared to microglia. Inhibition of NFκB or MAPK signaling revealed that 40Hz flicker-induced cytokines are mediated in part by NFκB and MAPK signaling, while 40Hz flicker-induced changes in microglial ramification were mediated by NFκB signaling only. Together, these results show that different flicker frequencies and flicker-induced brain rhythms elicit distinct microglia and cytokine responses mediated by the NFκB signaling pathway. These studies reveal novel effects of different brain rhythms in microglia and cytokines and important mediators underlying how brain rhythms and flicker stimulation rapidly elicits changes in brain signaling and cytokines.

Interactions between neurons, microglia, and cytokines support neuron health and survival and are key to normal brain functions, like development and learning (*14, 15, 54, 55*). Moreover, microglial dysfunction plays a key role in multiple diseases, for example by overly pruning synapses in anxiety and depression disorders or failing to clear pathogenic proteins that accumulate in neurodegenerative diseases (*13*). While prior studies have shown that broad increases or decreases in neural activity, like seizure activity or silencing, alter microglia, little work has shown how specific patterns of neural activity impact microglia (*15, 23, 56, 57*). This is especially important to understand because extensive research has shown that different frequencies of neural activity occur naturally and are associated with different brain functions (*58*). Our own recent work has shown that 40Hz frequency neural activity recruits microglia into an engulfing state in mouse models of Alzheimer’s disease (*7–9*). However, prior studies have primarily considered pathological states and 40Hz (gamma) activity. Whether these effects generalize and indicate that brain rhythms beyond gamma control microglia responses in normal adult brains was unknown. We previously showed that 40Hz flicker alters cytokine protein expression in healthy mice (*10*), which is consistent with our current findings that 40Hz flicker alters transcription factors in control of cytokine expression (**Fig. 3**). Here we also found that flicker-induced neural activity has distinct frequency-specific effects on microglia in healthy mice. We compared 40Hz flicker to 20Hz flicker based on our earlier findings that 20Hz flicker induces 20Hz neural activity without changing mean firing or behavior, and that 20Hz flicker leads to lower cytokine expression. Because microglial morphology changes as microglia shift between surveillance and engulfing states, we examined microglial morphology histologically after 40Hz and 20Hz flicker (*14, 38*). We found that 1hour of 40Hz or 20Hz flicker led to significant differences in microglial morphology in visual cortex. Specifically, after 20Hz flicker, microglia have longer, more branched processes closer to the cell soma with smaller cell bodies, changes that are consistent with surveillance functions, while 40Hz had the opposite effects (*14, 38*). The microglia morphological changes in response to 40Hz are similar to those we observed in prior studies in animal models with amyloid beta pathology, revealing that microglia responses to 40Hz flicker are not specific to Alzheimer’s disease pathology. Interestingly, microglia morphology with shortened processes and enlarged soma are typically observed during development or under pathological conditions when microglia engulf synapses or pathogens. Thus, we discovered a non-pathological stimulus that induces this microglia morphology in adulthood in healthy brains. Importantly, the current study was conducted in healthy mice in the absence of pathology, showing the effect of flicker on brain immunity is a general process that may play a role in normal brain function.

We have found that NFκB and MAPK phospho-signaling pathways mediate the effects of flicker-induced gamma on cytokines and microglia. Prior work studying the effects of neural activity on microglia has typically not examined NFκB and MAPK phospho-protein signaling pathways (*59*– *62*). Thus, this study reveals new mechanisms of neural activity-cytokine-microglia interactions. While the MAPK and NFκB pathways and downstream cytokine signaling represent canonical immune-regulatory pathways, they work in concert with diverse other pro- and anti-inflammatory pathways. These pathways are parallel to, or mediate the functions of, diverse other immune-regulatory mechanisms, including Jak/STAT (*63*), PI3K/Akt/mTOR (*64*), the complement system (*65*), sphingosine-1-phosphate (*66*), scavenger receptors (*67*), among many others. Moreover, microglia express purinergic receptors, including P2Y12, which can evoke migration and activation (*68*). Although we do not rule out the importance of any of these pathways, we did not interrogate them in the current study because of our prior findings that 40Hz flicker stimulates rapid MAPK and NFkB pathway activity within 5mins. Furthermore, our unbiased gene RNAseq screen and pathway analysis (**Fig. S3**) did not implicate these alternative mechanisms.

We found that flicker induced the expression of a few cytokines even when microglia were depleted. While microglia are the commonly accepted source of cytokines, prior evidence shows that neurons also release cytokines (*20–23*). Furthermore, we found that some cytokines are microglia-dependent. Surprisingly, however, some flicker-induced cytokines were microglia-independent including IL-10 and M-CSF (**Fig. 3**). M-CSF promotes microglia survival, proliferation, and phagocytosis, and IL-10 has anti-inflammatory functions (*69–74*). Therefore, these results establish that a subset of flicker induced cytokines do not arise from microglia. Because 40Hz flicker-induced cytokine expression requires NFκB and some of these cytokines are microglia-independent, we wondered from in what cell-types NFκB is activated. We found that pNFκB was highly colocalized with neurons following 40Hz flicker. These results show that neurons are abundant sources of pNFκB, which leads to cytokine expression and microglia recruitment in healthy adult mice and uncover a new role for NFκB in gamma frequency effects on microglia and cytokine signaling.

The present study has several limitations that suggest avenues of future research. First, we utilized small molecule MAPK (JNK+ERK) and NFκB signaling inhibitors to define the roles of these pathways in mediating cytokine expression in response to 40Hz flicker. We selected these inhibitors because they inhibit cell signaling across multiple cell types, but this leaves open the question of which cells play a causal role in the observed cytokine-suppressive effects. We believe this concern is partially abrogated because we quantified cytokines 1 hour after the start of stimulation, and thus secondary effects through other cell types would be unlikely to have sufficient time to translate into protein changes or would have minimal effects (*75*). Moreover, we have shown that phospho-NFκB co-localizes with neurons, meaning that the main effects of these phospho-signaling inhibitors may be in neurons. Nevertheless, future cell-type specific knockout studies will confirm whether neuronal signaling is indeed responsible for flicker-induced cytokine expression. Additionally, it is possible that NFκB inhibitor acts directly on microglia by inhibiting NFκB in microglia, thus flicker could induce NFκB in microglia thus altering microglia and the production of cytokines. Secondly, we used the small molecule CSF1R inhibitor PLX3397 to define the role of microglia in mediating cytokine expression in response to 40Hz flicker because it effectively depletes microglia without the need for transgenic animals or induction of Cre-recombination using tamoxifen, which is immunomodulatory (*76*). However, it has recently been shown that CSF1R inhibitors can have direct effects on neurons in addition to their well-known effects on peripheral monocytes (*77*). We are encouraged that several of our most important up-regulated cytokines (e.g. M-CSF, IL-10) in response to 40Hz flicker are preserved in in CSF1R-treated/microglial depleted mice. We therefore conclude that these cytokines are secreted independent of microglia. Future studies will verify the roles of microglia using Cre-Lox mediated microglial depletion. Furthermore, brain rhythms are diverse, not only with different frequencies of activity but also different circuit mechanisms and relationships to sensory stimuli. Thus further work is needed to fully elaborate how the diverse array of brain rhythms affect microglia and cytokines. Lastly, we conducted the present work in males to be consistent with prior studies, but the effects of brain rhythms on microglia and cytokine responses in females needs to be established.

In total, we reveal a novel causal link between flicker, brain rhythms, microglial state, and the secretion of cytokines via the NFκB pathway. Based on our combined results, we propose a model in which specific frequencies of neural activity, including gamma, increase activation of NFκB phospho-signaling in neurons, leading to neuronal cytokine release, which in turn alters microglial state. This model is in line with prior work showing that NFκB phospho-signaling in neurons is regulated by synaptic activity (*28–31*). Thus, our data reveal new mechanistic bases for frequency specific neuron-to-microglia signaling elicited by visual flicker and brain rhythms. This neuronal regulation of microglia and cytokine responses may play a role in plasticity, since gamma oscillations have been shown to increase during learning and memory, or in response to injury (*78*– *80*). Furthermore, because we find flicker effects generalize beyond the context of Alzheimer’s pathology, this sensory-induced and brain rhythm-induced regulation of microglia and cytokine responses could be harnessed to mitigate pathological neuroimmune activity present in multiple diseases.

### Materials and Methods Mice

The Georgia Tech Institutional Animal Care and Use Committee or the Emory Institutional Animal Care and Use Committee approved all animal work performed in this study. For experiments assessing microglia, cytokines, and phosphosignaling, adult (2 to 3-month-old) male C57BL/6J mice were obtained from The Jackson Laboratory. Mice were pair-housed and acclimated to the vivarium for at least 5 days before experiments began. Food and water were provided *ad libitum*. Experiments were performed during the light cycle and conducted at different times of day and with varying order of stimulus presentation to control for circadian effects. Animals were randomly assigned to flicker exposure groups and experimenters were blind to flicker exposure conditions during analysis for all experiments. For widefield imaging experiments, *EMX1*-Cre male mice were bred in-house to allow neonatal injection of the AAV vectors on P_0_ (described below). Mice for imaging experiments were housed on reverse light cycle (12 h light and 12 h dark), and imaging experiments were performed during the dark cycle. Mice started to receive water restriction 5 days prior to the experiments and were habituated to the head-fixation setup.

### Voltage Imaging of Visual Cortex

To achieve pan-cortical expression of voltage indicators and reference fluorescence for widefield imaging, P0 EMX1-Cre pups received ICV injections bilaterally with 4 uL of AAV vectors (2 uL/hemisphere) (59). The injected solution consisted of AAV.php.eb-EF1a-DIO-JEDI-1P-Kv2.1-WPRE (3 × 1012 vg/mL) and AAV9-hSyn-mCherry (9 × 1012 vg/mL). A 10-uL Nanofil syringe and a 34G beveled needle (World Precision Instrument) was used to load and inject the vector mixture. The injection site was identified using 2/5 of the distance between the lambda suture to each eye with a targeting depth of 3mm.

Mice (3-months old) that expressed voltage sensor, JEDI-1P-Kv, and reference fluorescence, mCherry, were implanted with an imaging window using the clear skull preparation (60, 61) and headplate and were allowed to recover for 7 days after surgery. Widefield voltage imaging was conducted using a high-speed CMOS imaging system (MiCAM-Ultima, SciMedia Ltd) at 200 fps. During voltage imaging experiments, a light barrier was placed around the imaging window to prevent visual flicker from contaminating imaging signals. Animals were imaged during 20Hz and 40Hz visual flicker for 10 trials in which each trial consisted of 10s baseline (no flicker) and 10s flicker stimulation. Light-emitting diodes (LED) were used to apply visual flicker with an intensity approximately 400 lux at the head of the animal.

### Voltage Imaging Analysis

Signals from 100×100 pixels (10mm × 10mm) were binned to groups of 2×2 pixels yielding 2500 (50×50) traces. A pre-processing pipeline was used to subtract background fluorescence, correct photobleaching, and regress hemodynamics, motion artifact, and potential light contamination caused by visual flicker [reference]. The JEDI-1P-kv signals were lowpass filtered at 70Hz. The reference mCherry signals were filtered at 0-1, 1-10, 10-19.5, 19.5-21.5, 20.5-38, and 38-42Hz respectively in order to regress out the shared non-voltage artifacts from the voltage signals using ordinary least squares methods. The extracted voltage signals were used for further analyses. To identify the cortical areas that responded to visual flicker, we calculated pixel-wise power spectrum. Pixel-wise power at 20Hz or 40Hz during either baseline or visual flicker was averaged, and the mean power during baseline was subtracted from that during visual flicker, n = 10 trials. The difference in power at the frequency of interest from binned pixels were used to generate spatial heat maps. The heat maps were resized to 100×100 pixels and masked with a cortical map adapted from Brain Explorer (Allen Institute for Brain Science, 2004) (*81*). The heat maps were used to select a 2×2-pixel region of interest (ROI) in V1 that responded strongly to visual flicker. Voltage signal at the selected ROI from each animal was used to calculate the power spectrum from each trial which was averaged to plot the mean power spectral density (PSD) during baseline and stimulation, respectively, using MATLAB DSP System toolbox (dsp.SpectrumAnalyzer).

### Visual Stimulation Exposure

Mice were habituated in a dark room in the laboratory for at least 1 hour prior to experimentation. At the start of experimentation, mice were transferred to enclosures for flicker exposure which had 3 opaque sides and one clear side that faced a strip of LEDs. Animals were exposed to either LED lights flashing at 40Hz frequency (50% duty cycle: 12.5ms light on, 12.5ms light off), 20Hz frequency (50% duty cycle: 25ms light on, 25ms light off), or no flicker for either 15 minutes or 1 hour. LED intensity was about 400 lux at the head of the animal. Immediately after stimulation exposure, mice were anesthetized with isoflurane, and within 3 minutes mice were decapitated and brains were removed. The left hemisphere’s visual cortex was micro-dissected, placed in microcentrifuge tube, and flash frozen using liquid nitrogen to be used for cytokine and phospho-protein assays. The right hemisphere was used for immunohistochemistry.

### Phospho-protein Inhibitors

The NFκB pathway was inhibited using IKK inhibitor (IMD0354 at 10mg/kg) combined with Curcumin (S1848 at 100mg/kg). The MAPK pathway was inhibited using MEK inhibitor (SL327 at 100mg/kg) and JNK inhibitor (SP600125 at 15mg/kg). All drugs were solubilized in vehicle consisting of dimethyl sulfoxide (DMSO), polyethylene glycol (PEG), and sterile physiologic saline. Drugs were delivered via intraperitoneal (IP) injection 30 minutes before the start of 1hour of flicker stimulation.

### Microglia Depletion

Microglia were depleted using Pexidartinib (PLX3397, MedChem) incorporated into Open Standard Diet with 15Kcal% fat (Research Diets, INC) at a dose of 290mg/kg (**Fig. S4**). A control group was fed a diet of the Open Standard Diet with 15Kcal%. Mice were fed depletion or control diet for three weeks, and animal weight was monitored to ensure weight gain or loss did not differ between groups. Microglia depletion was confirmed via histology.

### Immunohistochemistry (IHC) and Microscopy

Following brain extraction, right hemispheres were drop fixed into cold 4% paraformaldehyde (PFA) for 24 hours, rinsed in 1X Phosphate Buffered Saline (PBS), and stored in PBS with 0.02% sodium azide (NaN_3_). Each hemisphere was rinsed and placed in 30% sucrose for three days and then frozen. Sagittal sections were obtained at 30 microns for phosphoprotein staining and 40 microns for microglia staining. Two sections per animal were rinsed 3 times for 10 minutes each in 1XPBS on a shaker, blocked for 1 hour in blocking buffer (0.2% Triton X, 5% Normal Goat/Donkey Serum, 1XPBS), then incubated overnight at 4°C in primary antibody (anti-pNFκB, anti-NeuN, anti-IBA1) diluted in blocking buffer at antibody combination and concentrations shown in **Table S2**. Sections were then washed in 1XPBS 3 times for 10 minutes each and incubated on shaker at room temperature for 2 hours in secondary antibody solution (goat anti-rabbit 555, goat anti-mouse 488) diluted in blocking buffer with antibody combinations and dilutions shown in **Table S2**. Sections were then washed in 1XPBS 3 times for 10 minutes each, and nuclei were stained with 4′,6-diamidino-2-phenylindole (DAPI) for 1 minute. Sections were mounted using Vectashield and Z-stack images of visual cortex were taken on a Nikon Crest Spinning Disk Confocal Microscope with a 40x objective or a Zeiss LSM700 with a 20x objective for glial imaging and reconstruction and pNFκB colocalization, respectively. For pNFκB colocalization, non-specific background staining was reduced by increasing the salt concentration (NaCl) in the blocking buffer from 0.3 to 0.5 M (*82*)

### Cytokine and Phospho-protein Assays

For signaling and cytokine analysis, the visual cortex was thawed on ice and lysed using Bio-Plex lysis buffer (Bio-Rad). After lysing, samples were centrifuged at 4 °C for 10 minutes at 13,000 RPM. Protein concentrations in each sample were determined using a Pierce BCA Protein Assay (Thermo Fisher). Total protein concentrations were normalized in each sample using Bio-Plex lysis buffer (Bio-Rad) and 6μg was loaded for cytokine analysis. Cytokine analysis was conducted using the Milliplex Mouse MAP Mouse Cytokine/Chemokine Magnetic Bead Panel 32-Plex Kit (Eotaxin, G-CSF, GM-CSF, IFN-γ, IL-1α, IL-1β, IL-2, IL-3, IL-4, IL-5, IL-6, IL-7, IL-9, IL-10, IL-12p40, IL-12p70, IL-13, IL-15, IL-17, IP-10, KC, LIF, LIX, MCP-1, M-CSF, MIG, MIP-1α, MIP-1β, MIP-2, RANTES, TNF-α, and VEGF). All kits were read on a MAGPIX system (Luminex, Austin, TX).

### Morphology and Colocalization Analysis

Bitplane Imaris 9.7.1 (Oxford Instruments, Concord, MA) was used to assess morphology of IBA-1+ microglia and colocalization of pNFκB with both NeuN+ neurons and IBA-1+ microglia while blinded to treatment groups. Surface reconstruction parameters were optimized across samples and examined by eye while blinded and prior to any hypothesis testing. To assess microglia morphology, only microglia whose processes were completely within the bounds of the image were included in analysis. Z-stack images of IBA-1+ stained visual cortex tissue were modified using Gaussian Filtering and cropped to standard dimensions (2301×3301×101 μm). Within Imaris, microglia were automatically reconstructed as surfaces utilizing standardized thresholds pertaining to diameter of clustered signal and level of surface detail. Irrelevant surfaces were filtered by number of voxels and total area. Surfaces were then manually corrected based on visual inspection. Surfaces that belonged to the same cell were grouped together as a single surface. Following reconstruction, microglia surfaces were filtered again to remove any remaining surfaces not part of a reconstructed cell. Surfaces were transformed into processes based on standardized thresholds including diameter, gaps between surfaces, and surface detail, and process properties were quantified. We measured soma area, total process length, branching depth, and number of branches at each Sholl radii from the soma (**Fig. S7**).

To assess colocalization of stained pNFκB with both NeuN+ neurons and IBA-1+ microglia, Z-stack images of stained visual cortex tissue were modified using baseline subtraction and gamma correction filtering and cropped to standard dimensions (2400 × 2400 × 120 μm). Within Imaris, cells (either neurons or microglia) were reconstructed as surfaces utilizing standardized thresholds pertaining to diameter of clustered signal and level of surface detail. pNFκB staining was reconstructed as surfaces using a different set of thresholds. Using both sets of surfaces, all pNFκB staining was eliminated except for the surfaces which were located a maximum distance of 5 μm from the reconstructed cells. Volumetric percent of total pNFκB compared to overlap of pNFκB staining with NeuN or IBA-1 cell stain was used to determine if pNFκB colocalized to a cell type.

### Partial Least Squares Discriminant Analysis

Data was z-scored within each group, and a partial least squares discriminant analysis (PLSDA) was performed in MATLAB (Mathworks) using the algorithm by Cleiton Nunes (Mathworks File Exchange). To identify latent variables (LVs) that best separated conditions, an orthogonal rotation in the plane of the first two latent variables (LV1-LV2 plane) was performed. Error bars for LV1 figures represent the mean and standard deviations after iteratively excluding single samples (a leave-one-out cross validation) and recalculating the PLSDA 1,000 times. To test different flicker conditions and durations, cytokine experiments were performed over several cohorts and samples were normalized by removing batch effects. We performed a batch effects analysis (combat package in R v 3.5.0) to remove any between-experiment variability before conducting the PLSDA. Lastly, for analyzing the effect of 40Hz flicker following microglia depletion, the data from three cohorts was combined by performing a batch-effects analysis (combat package in R version 3.2.0) to remove any between-experiment variability prior to conducting the PLSDA.

### RNA Sequencing

mRNA were quantified from male 4–6-month-old WT mice exposed to either 20Hz or 40Hz audio/visual flickering for 1h (n=8). RNA was isolated from visual cortices using a Qiagen RNeasy kit (217804; Qiagen; Hilden, Germany) according to the manufacturer’s protocol. Libraries were sequenced for paired-end mRNA using the NovaSeq 6000 Sequencing System to obtain a sequencing depth of 30-40 million reads per sample. Prior to sequencing, quality control was performed using a bioanalyzer for RNA Integrity Number (RIN) greater than 7. A NEBNext Poly(A) mRNA Magnetic Isolation Module and NEBNext Ultra II Directional RNA Library Prep Kit was used to prepare sequencing libraries. RNA alignment was performed by Molecular Research Lab and data were validated for fastq integrity and quality. Specifically, each sample was run on four lanes and merged, followed by using DNAstar ArrayStar. Next, Qseq reads were mapped using the mouse reference genome GRCm39. For read assignment, the threshold was set at 20bp and 80% of the bases matching within each read. Duplicated reads were eliminated, genes with less than 20 total raw counts were removed from analysis. All gene counts were normalized using the DESeq2 package in R. RNA sequencing data are deposited on NCBI gene expression omnibus.

### Gene Set Variation Analysis (GSVA) and Statistical Analysis of Gene Expression Data

To study holistic gene changes within different pathways relevant to brain immunity and neurotransmission function, GSVA analysis was conducted. GSVA is an unsupervised enrichment algorithm which identifies variations of pathway activity by defining enrichment score for gene sets which each contain a set of genes with shared cellular function. For GSVA gene set reference and symbols, the Molecular Signatures Database C2 [v7] gene sets (MSigDB) were used. To compare statistical differences between groups for each pathway of interest, enrichment scores for each gene set between groups were computed by measuring the true differences in means versus the differences computed by a random distribution obtained by permuting the gene labels 1000 times. False discovery rate (FDR) adjusted p-values were computed for detection of differences between two groups and significant level was set at FDR < 0.25. All data is presented as mean ± SEM. Non-parametric Wilcoxon rank sum test was used to determine the group differences (40Hz vs. 20Hz) in individual genes within each pathway. p < 0.05 was considered statistically significant. All analyses were performed in R using the *stats* package v.3.6.2.

### Statistical Analysis

For morphology and colocalization analyses, hypothesis testing was performed between the four groups using one-way, two-tailed, unpaired ANOVAs assuming equal variance, with an alpha-value of 0.05. Differences between groups were found from post-hoc multiple-comparisons tests with a Tukey’s HSD correction. We used the mixed-model procedure in SPSS 26 (IBM) for Sholl analyses with fixed-effects for the intercept, frequency, and radii, identity covariance structure, and maximum-likelihood estimation. For multi-plex analysis, either a one-way ANOVA (more than two groups) or two-tailed unpaired t-test (two groups) was used to determine if there was a significant LV1 separation between groups. The top correlated cytokines and/or phospho-proteins of LV1 were isolated and an ANOVA or two-tailed unpaired t-test was performed using GraphPad Prism 8 (GraphPad Software, La Jolla, CA) to determine statistical significance between groups. These tests were followed by a post-hoc Dunnett’s multiple comparisons test to determine differences between specific groups or a Tukey’s multiple comparisons test to compare differences in phosphorylation levels across time. To further confirm significant differences between groups using PLSDA, a permutation analysis was conducted, which randomly assigned animals into experimental groups and ran the PLSDA based on these shuffled values 1000 times. The mean for each permuted value was then compared as the square root of the difference between the sums of squares on the LV1 and LV2 axis of PLSDA. For each test, true group assignment showed p_permute_<0.05 compared to the randomly permuted distribution, further confirming the validity of our data. To compare individual cytokine expression in the microglia depletion and flicker experiment, a one-way ANOVA was used, following by multiple comparisons using the Tukey method.

## Acknowledgments

We thank the Wood and Singer labs for critical feedback and technical assistance on this work.

## Funding

National Institutes of Health grant R01-NS-109226 (ACS)

National Institutes of Health grant R01-NS-109226-01S1 (KMG)

National Institutes of Health grant R01-NS-111470 (DJ)

The Coins for Alzheimer’s Research Trust Fund (LBW/ACS)

Packard Award in Science and Engineering (to A.C.S.)

Friends and Alumni of Georgia Tech (A.C.S.)

NSF CAREER 1944053 (LBW)

## Author contributions

Conceptualization: KMG, YW, DJ, LBW, ACS,

Methodology: KMG, AMP, AS, CH, SB, YW, MG, DJ, LBW, ACS

Investigation: KMG, AMP, AS, CH, SB, YW, MG, DJ, LBW, ACS

Visualization: KMG, AMP, AS, CH, SB, YW, MG, DJ, LBW, ACS

Supervision: DJ, LBW, ACS Writing—original draft: AMP, AS, LBW, ACS

Writing—review & editing: AMP, AS, CH, SB, LBW, ACS

## Data and materials availability

All data and code will be made available at the time of publication via GitHub for custom code, NCBI GEO for transcriptomic data, ProteomeXchange (PX) consortium of proteomics resources for proteomic data, NeuroMorpho.Org for histological data.

