## Supplementary Materials for "Brain rhythms control microglial response and cytokine expression via NFκB signaling"

**This PDF file includes:**

Figs. S1 to S8
Tables S1
Caption for Movie S1
Caption for Data S1

**Other Supplementary Materials for this manuscript include the following:**

Movie S1
Data S1

**Supplementary Figures**

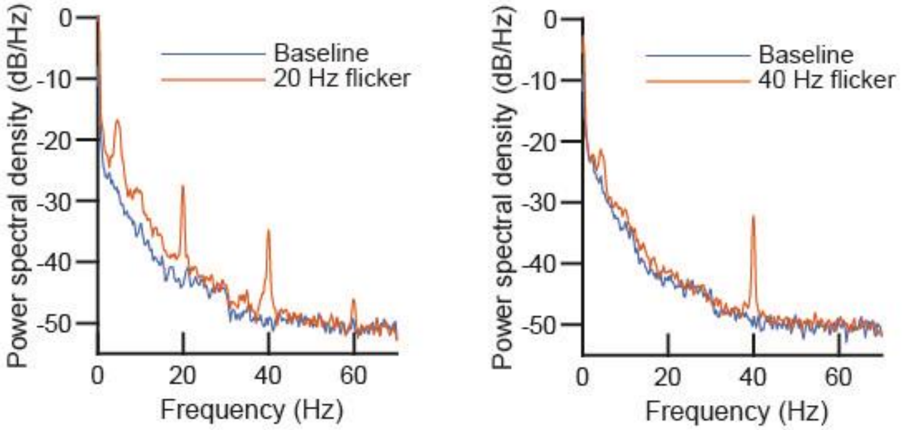


**Fig. S1. 40Hz visual flicker stimulates primary visual cortex. Left:** Mean power spectral density (PSD) of voltage imaging signal (2×2-pixel) at primary visual cortex during 20 Hz visual flicker (orange) and baseline (blue) in EMX1-Cre mice expressing JEDI-1P-kv and mCherry. *Note peaks at 40Hz and 60Hz are expected harmonics of 20Hz due to the fact that the neural response is not a perfect sinusoid. **Right:** Mean PSD during 40 Hz visual flicker. Shaded areas denote 95% CI. n = 6 mice, 60 trials, 10 trials/mouse.

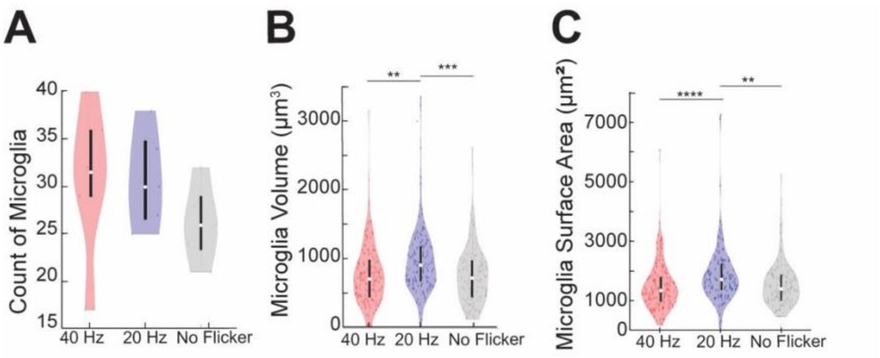


**Fig. S2. Visual flicker affects microglia volume and surface area in a frequency specific manner in WT mice.** Analysis of IBA-1+ microglia after animals were exposed to 1hr of 40 Hz Flicker (red, n=6 mice, 174 microglia), 20 Hz Flicker (blue, n=5 mice, 140 microglia), or No Flicker (grey, n=5 mice, 132 microglia) stimulation for **(A)**, number of microglia (F=0.9797, p=0.4015) (F(2,13)=0.9797, p=0.4015; post-hoc t-tests: 40Hz vs. 20Hz: p = 0.9999, 40Hz vs No Flicker: p = 0.446, 20Hz vs No Flicker: p = 0.47997; Tukey's HSD corrected for multiple comparisons) **(B)**, Volume of IBA-1+ microglia (including cell body and processes) after 40Hz, 20Hz, or No Flicker stimulation (F(2,443)= 9.9694, p=0.0001; post-hoc t-tests: 40Hz vs. 20Hz: p = 0.0002, 40Hz vs No Flicker: p = 0.9957, 20Hz vs No Flicker: p = 0.0004; Tukey's HSD corrected for multiple comparisons) and **(C)**, surface area of whole IBA-1+ microglia after 40Hz, 20Hz, or No Flicker stimulation (F(2,443)= 9.4815, p=9.39E-5; post-hoc t-tests: 40Hz vs. 20Hz: p = 9.39E-5, 40Hz vs No Flicker: p = 0.7809, 20Hz vs No Flicker: p = 0.0035; Tukey's HSD corrected for multiple comparisons). Box plots inside violin plots indicate median and quartiles, dots indicate individual microglia. F- and p-values were generated from one-way, two-tailed, unpaired ANOVA tests. Differences between groups were found from one-way, two-tailed, t-tests with Bonferroni correction. **p<0.01,***p<0.001,****p<0.0001.

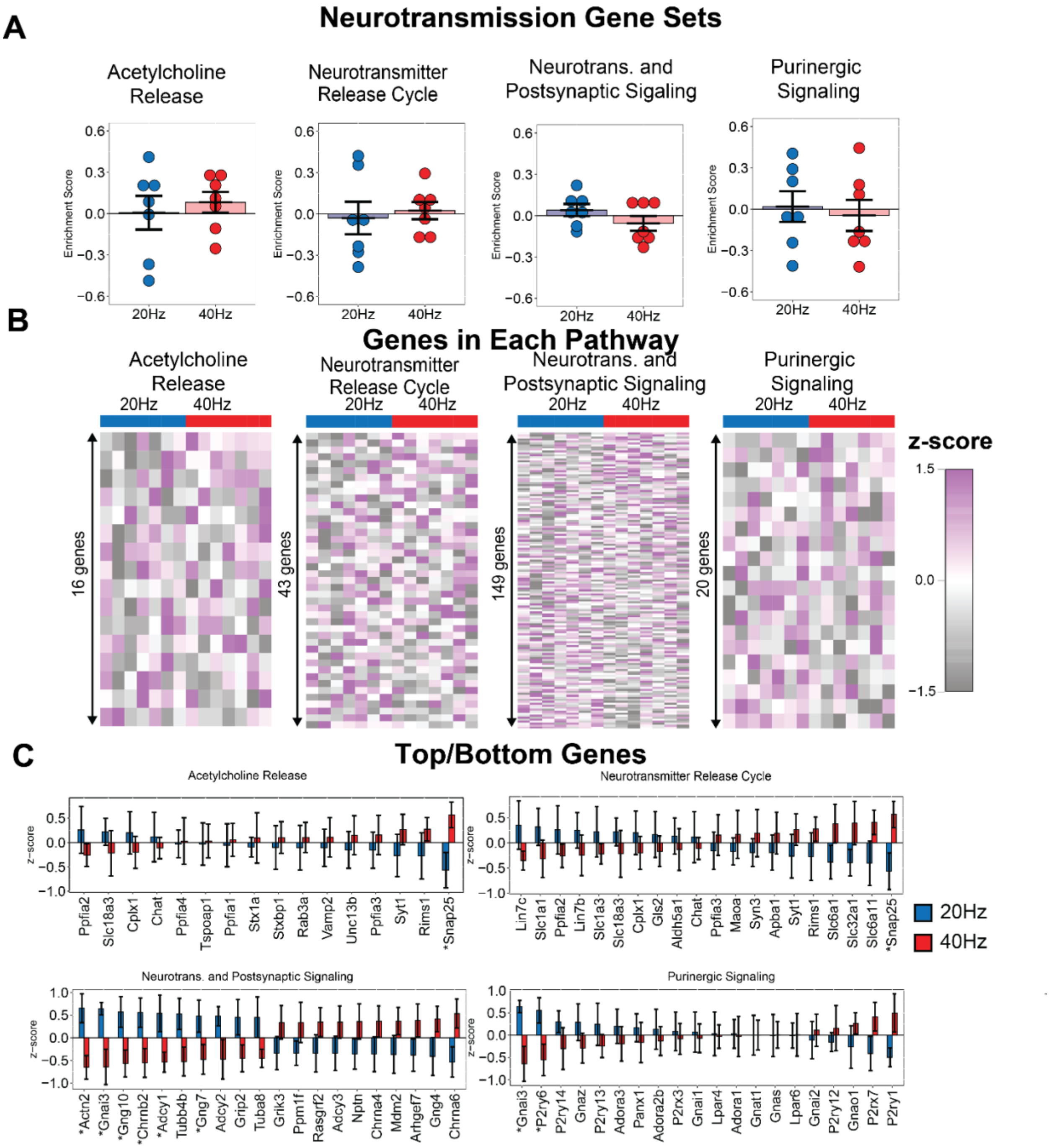


**Fig. S3. Flicker audio/visual stimulation for 1 hr at 40 Hz and 20 Hz does not significantly modulate gene sets associated with neurotransmission.** (**A**) Gene set variation analysis did not identify significant enrichment immune-related gene sets (mean±SEM, statistical testing via gene set permutation analysis, dots indicate individual animals). (**B**) Heatmap representation of all genes within each pathway (rows are z-scored). (**C**) Few genes within neurotransmission-related pathways are individually significantly different. (DESeq2, *p<0.05 un-adjusted).

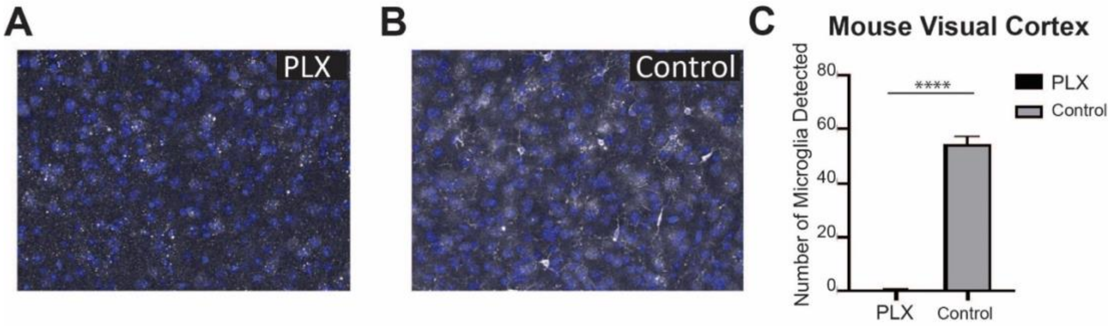


**Fig. S4. PLX 3397 successfully depletes microglia while cytokines remain expressed.** Microglia were depleted using Pexidartinib (PLX3397, MedChem) incorporated into the Open Standard Diet with 15Kcal% fat (Research Diets, INC) at a dose of 290mg/kg for 3 weeks. A control group was fed a diet of the Open Standard Diet with 15Kcal%. (A) Representative immunohistochemistry for IBA1+ microglia in the mouse visual cortex following 3-weeks of PLX diet compared to (B) mice fed control chow. (C) Number of microglia in visual cortex in mice fed PLX (black) and control (grey) diets ($t(8)=17.35$, $p<0.000001$). Error bars indicate mean±SEM.

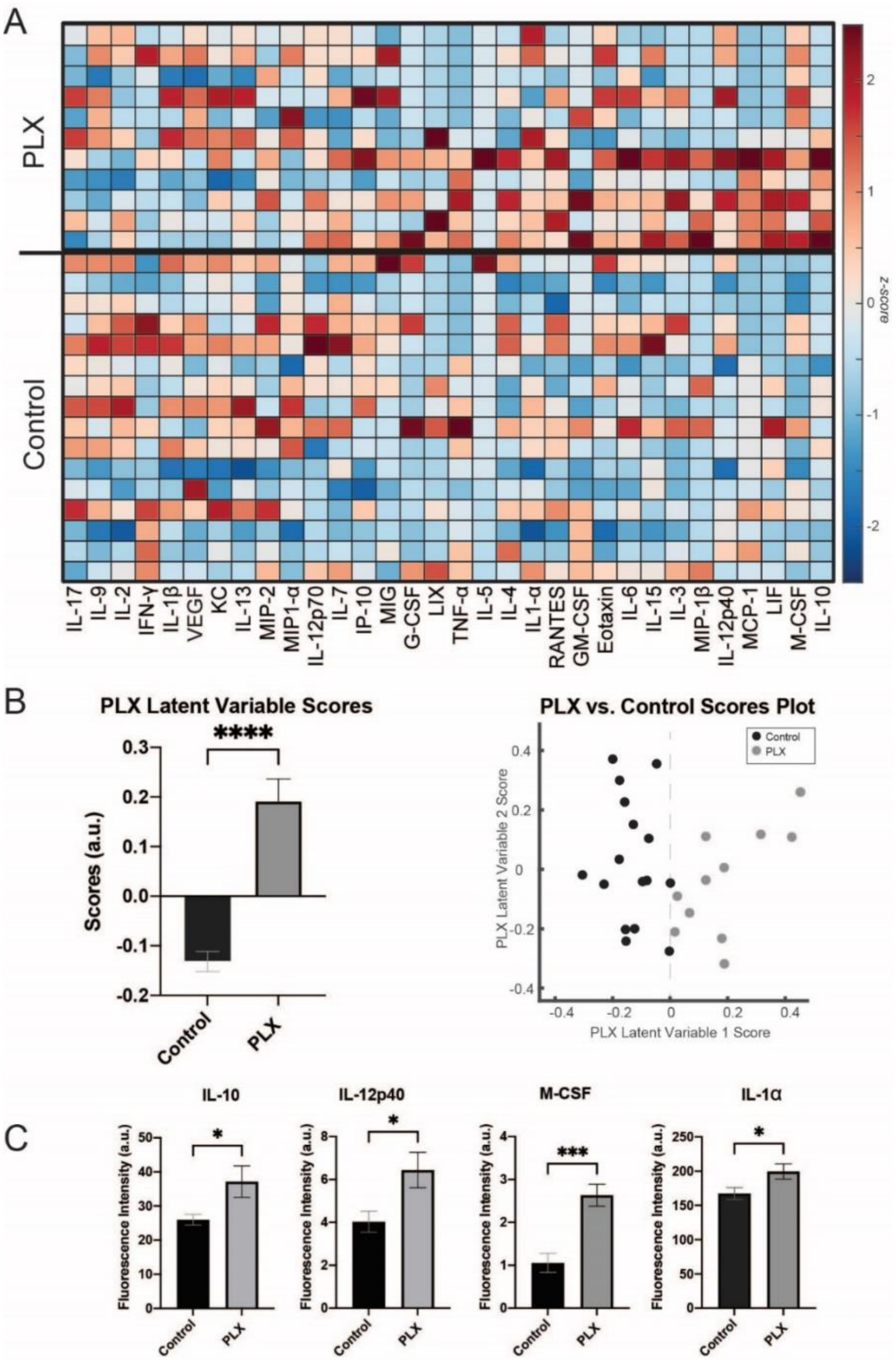


**Fig. S5. Cytokines Persist in the Absence of Microglia.**

**(A)** Cytokine expression in visual cortices of mice fed PLX3397 or control diet for three weeks. Each row represents one animal. Cytokines (columns) are arranged in the order of their weights on the LV1. Color indicates z-scored expression levels for each cytokine. **(B)** (**right**) PLSDA identified LV1, the axis that separated animals fed PLX3397 and a control diet, (**left**) LV1 scores were significantly different between the two groups of animals (mean ± SEM; $t(25)=7.297$, $p<0.0001$, unpaired, two-tailed t-test). **(C)**, Cytokines with significant differences in expression between both groups, IL-10: $t(25)=2.647$, $p=0.0139$; , IL-12p40: $t(25)=2.665$, $p=0.0133$; M-CSF: $t(25)=4.608$, $p=0.0001$; IL-1α: $t(25)=2.309$, $p=0.0295$.SEM, standard error of the mean.

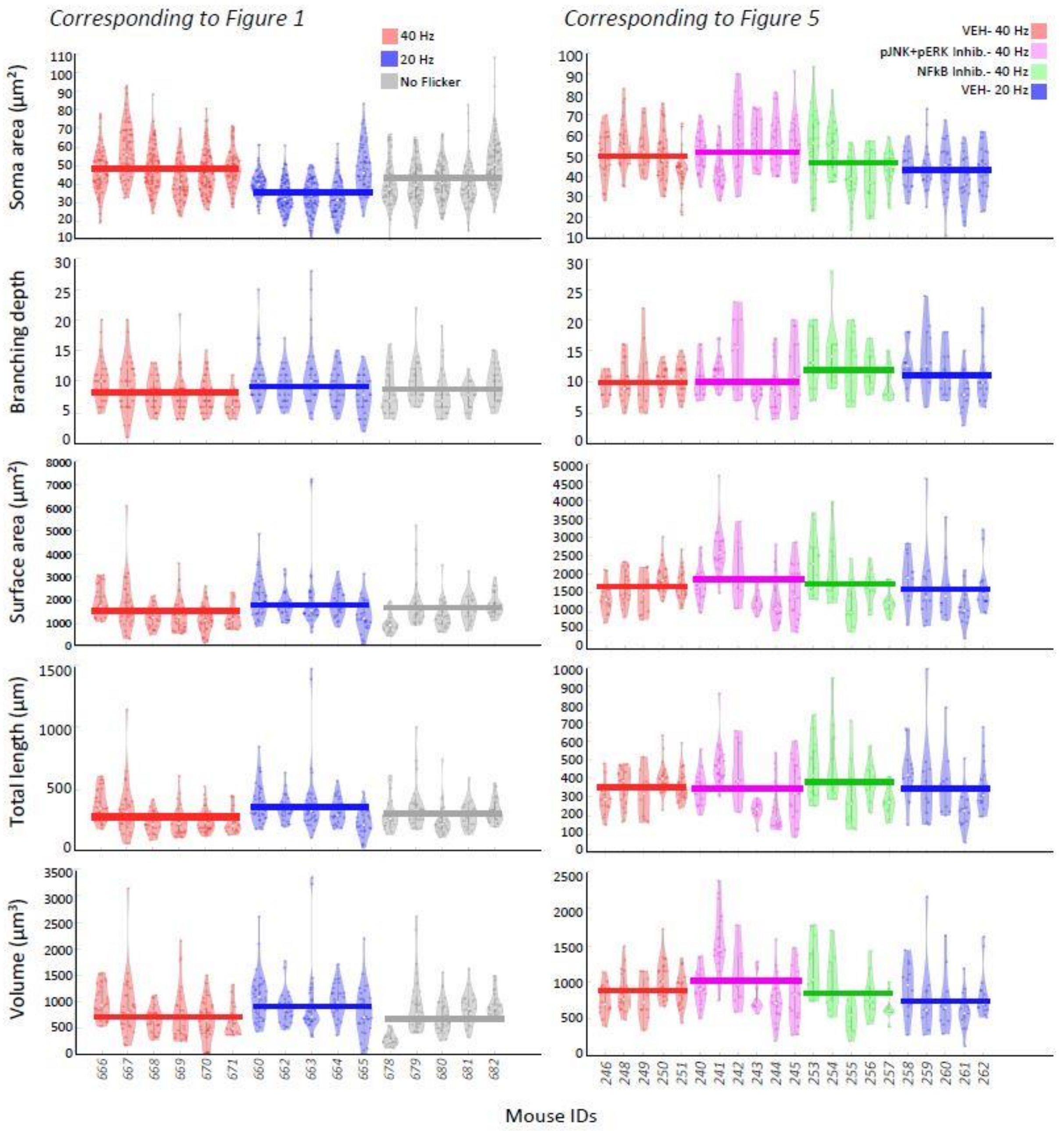


**Fig. S6. Microglia morphology measures per animal. Left) As in Fig 1 & Fig S2,** Analysis of IBA-1+ microglia after animals were exposed to 1 hour of 40 Hz Flicker (red, n=6 mice, 174 microglia), 20 Hz Flicker (blue, n=5 mice, 140 microglia), or No Flicker (grey, n=5 mice, 132 microglia) stimulation for soma area ($\mu m^2$), branching depth, whole-cell surface area ($\mu m^2$), total process length (µm), and total microglia volume ($\mu m^2$). **Right) As in Fig 5 & Fig S5** Analysis of IBA-1+ microglia measures as in the left column, after animals were exposed to one hour of 40 Hz Flicker and Vehicle Injection (orange, n=5 mice, 128 microglia), 40 Hz Flicker and pJNK+pERK Inhibitor injection (pink, n=6 mice, 129 microglia), 40 Hz Flicker and NFkB Inhibitor injection (green, n=5 mice, 116 microglia), and 20 Hz Flicker and Vehicle injection (blue, n=5 mice, 106 microglia). X-axes represent individual mice IDs. Dark colored line indicated mean per group.

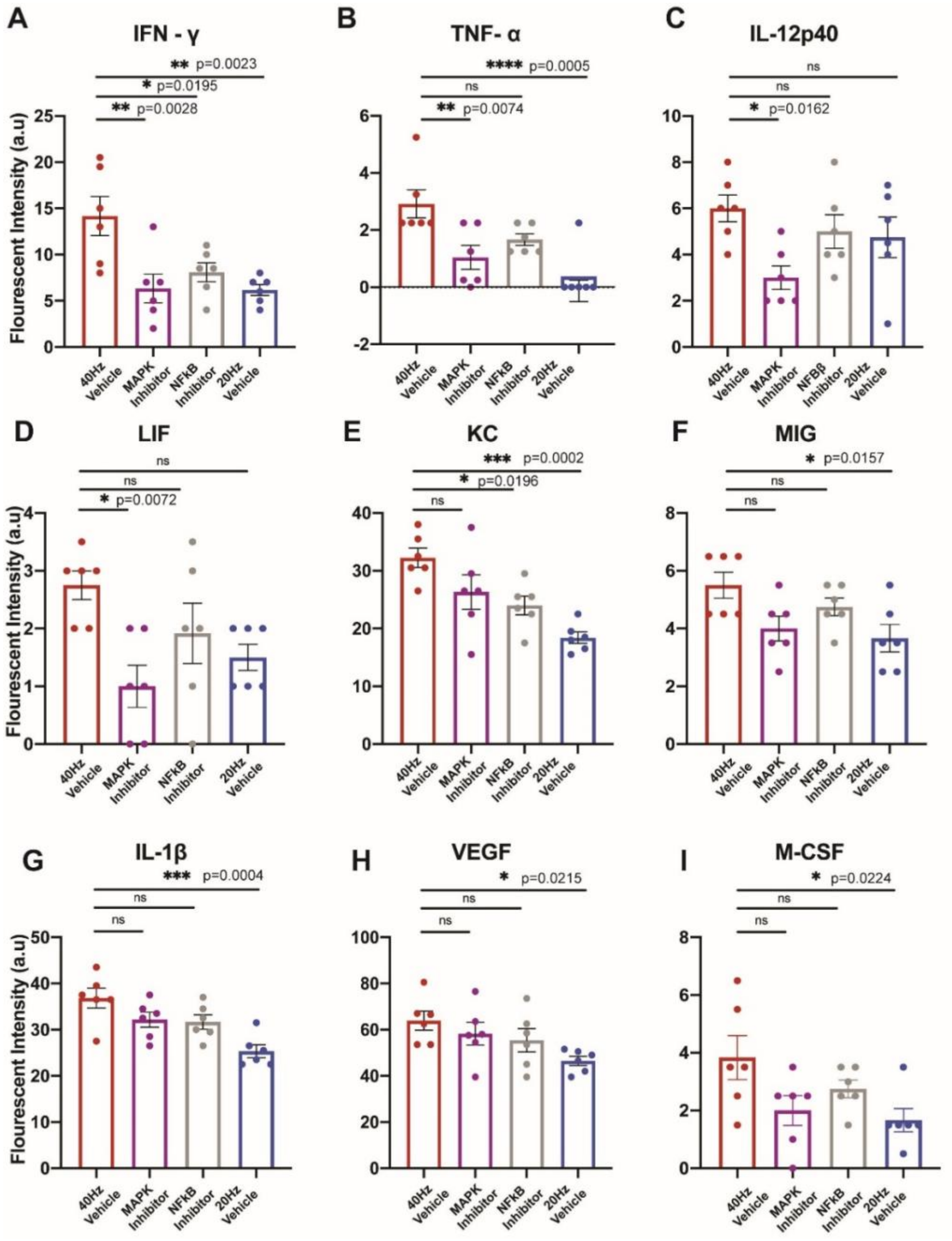


**Fig. S7. Inhibition of phosphoprotein pathways impact on key inflammatory cytokines. (A)** IFN-γ expression across groups, (F(3,20) = 6.787, p =0.0024, one-way ANOVA). The p-values from a Dunnett's multiple-comparison test are listed. **(B)**, As in A for TNF-α, (F(3,20) = 7.774, p =0.0012, one-way ANOVA). **(C)**, As in A for IL-12p40, (F(3,20) = 3.257, p =0.0431, one-way ANOVA). **(D)**, As in A for LIF, (F(3,20) = 4.225, p =0.0182, one-way ANOVA). **(E)**, As in A for KC, (F(3,20) = 8.57, p =0.0007, one-way ANOVA). **(F)**, As in A for MIG, (F(3,20) =3.782 , p =0.0268, one-way ANOVA). **(G)**, As in A for IL-1β, (F(3,20) =7.633 , p =0.0014, one-way ANOVA). **(H)**, As in A for VEGF, (F(3,20) =3.007, p =0.0545, one-way ANOVA). **(I)**, As in A for M-CSF, (F(3,20) =3.352 , p =0.0395, one-way ANOVA). Error bars indicate mean ± SEM, dots indicate individual animals.

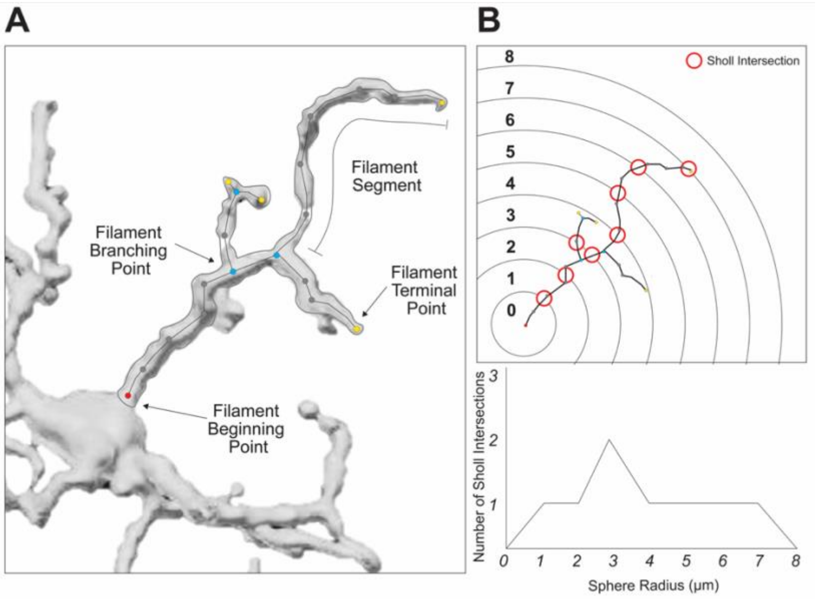


**Fig. S8. Microglia reconstruction protocol** (**A**) Diagram showing key features of filament segments superimposed on reconstructed microglia, such as filament beginning, terminal, and branching points. (**B**) Diagram demonstrating Sholl radii using a simplified microglia (*top*) and accompanying graph plotting number of Sholl intersections at each radius (*bottom*).

### Supplementary Tables

| | Primary | Secondary |
|---|---|---|
| Set 1 | pNFκB rabbit (ab86299) (1:500)<br>NeuN mouse (ab104224) (1:500) | Goat anti-rabbit 555 (A32732) (1:2000)<br>Goat anti-mouse 488 (A11001 (1:2000) |
| Set 2 | pNFκB rabbit (ab86299) (1:500)<br>GFAP mouse (MOB064-05) (1:500) | Goat anti-rabbit 555 (A32732) (1:2000)<br>Goat anti-mouse 488 (A11001 (1:2000) |
| Set 3 | pNFκB rabbit (ab86299) (1:500)<br>IBA1 goat (ab5076) (1:100) | Donkey anti-rabbit 488 (A21206) (1:2000)<br>Donkey anti-goat 555 (A21432) (1:2000) |

**Table S1. Antibodies and dilutions used for histology.** Primary and secondary antibodies used to visualize pNFκB and NeuN (**top**), pNFκB and GFAP (**center**), and pNFκB and IBA1 (**bottom**).

### Supplementary Movies

**Movie S1. (separate file)**
Mouse exposure to 40Hz visual flicker.

### Supplementary Data

**Data S1. (separate file)**

Statistical analysis (DESeq2) of individual genes within the pathways highlighted in Figures 2 and S3: Cytokine Signaling, Complement, Cellular Stress, MAPK Pathway, Acetylcholine Release, Neurotransmitter Release Cycle, Neurotransmitter and Postsynaptic Signaling, Purinergic Signaling.
